# Site-Specific Nanobody Inhibitors of the Proteasomal Deubiquitinase UCH37

**DOI:** 10.1101/2025.08.11.669711

**Authors:** Yanfeng Li, Lin Hui Chang, Rishi Patel, Elizaveta I. Shestoperova, Jiale Du, Mohamed Salam, Andrew N. R. Doig, Chittaranjan Das, Eric R. Strieter

## Abstract

Dysregulation of the ubiquitin (Ub) proteasome system (UPS) is linked to numerous human diseases, making its components attractive therapeutic targets. Among these, the proteasomal deubiquitinase UCH37 (also known as UCHL5) has two distinct Ub-processing activities: C-terminal Ub hydrolysis and K48-linkage–specific chain debranching. Separating these activities has not been possible with existing small-molecule inhibitors, which target the catalytic site and broadly affect both functions along with many other DUBs. Here, we report the development of two site-specific nanobody inhibitors of UCH37 that together enable this separation. Using yeast surface display, we generated Nb-cS1, which binds the canonical Ub-binding (S1) site and blocks both activities, and Nb-ncS1, which binds the non-canonical K48-specific (ncS1) site and selectively impairs chain debranching without affecting C-terminal Ub hydrolysis. X-ray crystallography and hydrogen-deuterium exchange mass spectrometry define the distinct binding surfaces engaged by each nanobody. In cells, both nanobodies selectively engage UCH37, with Nb-ncS1 showing the cleanest target engagement. These tools provide the first means to independently interrogate UCH37’s two enzymatic activities and offer new approaches for dissecting the distinct roles of UCH37 in proteasomal and chromatin-associated contexts.

## INTRODUCTION

The ubiquitin-proteasome system (UPS) plays a fundamental role in human biology by selectively targeting proteins for degradation^1^. It is a critical regulator of key processes such as the cell cycle^2^, transcription^3^, DNA repair^4^, and cellular responses to growth factors and stress^5,6^. Misregulation of the UPS is a hallmark of numerous human diseases, driving significant interest in developing therapeutic strategies to modulate its components^7–9^. Among these, deubiquitinases (DUBs)—the enzymes responsible for removing and editing ubiquitin modifications—are particularly attractive therapeutic targets due to their diversity and well-defined catalytic sites^10,11^. However, developing selective DUB inhibitors is often challenging, hindered by a lack of understanding of DUB biology and targets.

The challenge of developing selective inhibitors is especially relevant to the proteasomal deubiquitinase (DUB) UCHL5, also known as UCH37. UCH37 is a ubiquitously expressed DUB that is recruited to the distal portion of the 26S proteasome by the ubiquitin-binding subunit ADRM1/RPN13, and it also functions as a subunit of the INO80 chromatin remodeling complex^12–16^. Overexpression of UCH37 has been observed in several human cancers, including esophageal^17^, ovarian^18^, liver^19^, colorectal^20^, and gastric cancers^21^. In some cases, elevated levels of UCH37 have been linked to poor patient outcomes. However, the precise roles of UCH37 in the proteasome and the INO80 complex remain poorly understood, leaving the molecular basis for its association with pathogenesis unclear. Small-molecule inhibitors such as b-AP15 and VLX1570, based on a reactive α,β-unsaturated carbonyl motif, were developed primarily to target proteasome-associated deubiquitinases, including UCH37 and USP14, and have been evaluated in clinical trials^21–24^. Unfortunately, these molecules target a broad range of proteins, inducing nonspecific aggregation and general cellular toxicity^25,26^. As a result, the therapeutic potential of targeting UCH37 remains uncertain, and any conclusions drawn from using these molecules should be approached with caution.

We set out to develop a new generation of high-precision tools to interrogate UCH37 function, building on recent advances in its biochemical and structural characterization. Our lab, along with others, discovered that UCH37 not only removes small C-terminal fragments from ubiquitin (Ub) but also preferentially cleaves branched Ub chains containing K48 linkages, thereby promoting proteasomal degradation^27,28^. Structural studies revealed that UCH37 employs an aromatic-rich motif, composed of phenylalanines F117 and F121 on the face opposite the canonical ubiquitin-binding site, to specifically recognize K48-linked ubiquitin chains and remove branchpoints^29^. The spatial separation of the canonical S1 site from this non-canonical K48-specific (ncS1) site suggested that each activity might be inhibited independently with a sufficiently selective binder. Here, we report the development of two nanobodies that realize this possibility: Nb-cS1, which binds the canonical S1 site and blocks both C-terminal Ub hydrolysis and chain debranching, and Nb-ncS1, which binds the ncS1 site and selectively impairs debranching while leaving C-terminal hydrolysis intact. X-ray crystallography, hydrogen-deuterium exchange mass spectrometry, and cellular target-engagement assays establish the binding modes and selectivity of both nanobodies, and together they constitute the first tools capable of separating UCH37’s two enzymatic activities in biochemical and cellular settings.

## RESULTS

### Generation of Nanobodies Targeting UCH37

We employed a yeast surface-display library of synthetic nanobodies to screen for site-specific binders of UCH37 (Figure 1a, Supplementary Figures 1a-e)^30^. Our initial focus was on identifying nanobodies that target the K48 ubiquitin (Ub) chain-specific binding site, responsible for UCH37’s debranching activity. We hypothesized that blocking UCH37’s canonical S1 Ub binding site would increase the likelihood of isolating nanobodies that bind to the aromatic-rich motif comprising the K48-specific site (Figure 1a). To achieve this, we conjugated Ub propargylamine (Ub-PA) to the active site cysteine of UCH37 in the presence of the DEUBAD domain of RPN13, its proteasomal binding partner^31,32^. The DEUBAD domain is known to prevent UCH37 dimerization and enhance its debranching activity^33^. The resulting Ub-UCH37•RPN13^DEUBAD^ complex was then exposed to the yeast display library (Figure 1c). Magnetic-activated cell sorting (MACS) was used to remove non-binders, followed by fluorescence-activated cell sorting (FACS) to identify cells expressing high levels of nanobodies (c-myc tag) and robust binding to Ub-UCH37•RPN13^DEUBAD^ (biotin tag) (Figure 1d).

**Figure 1:**
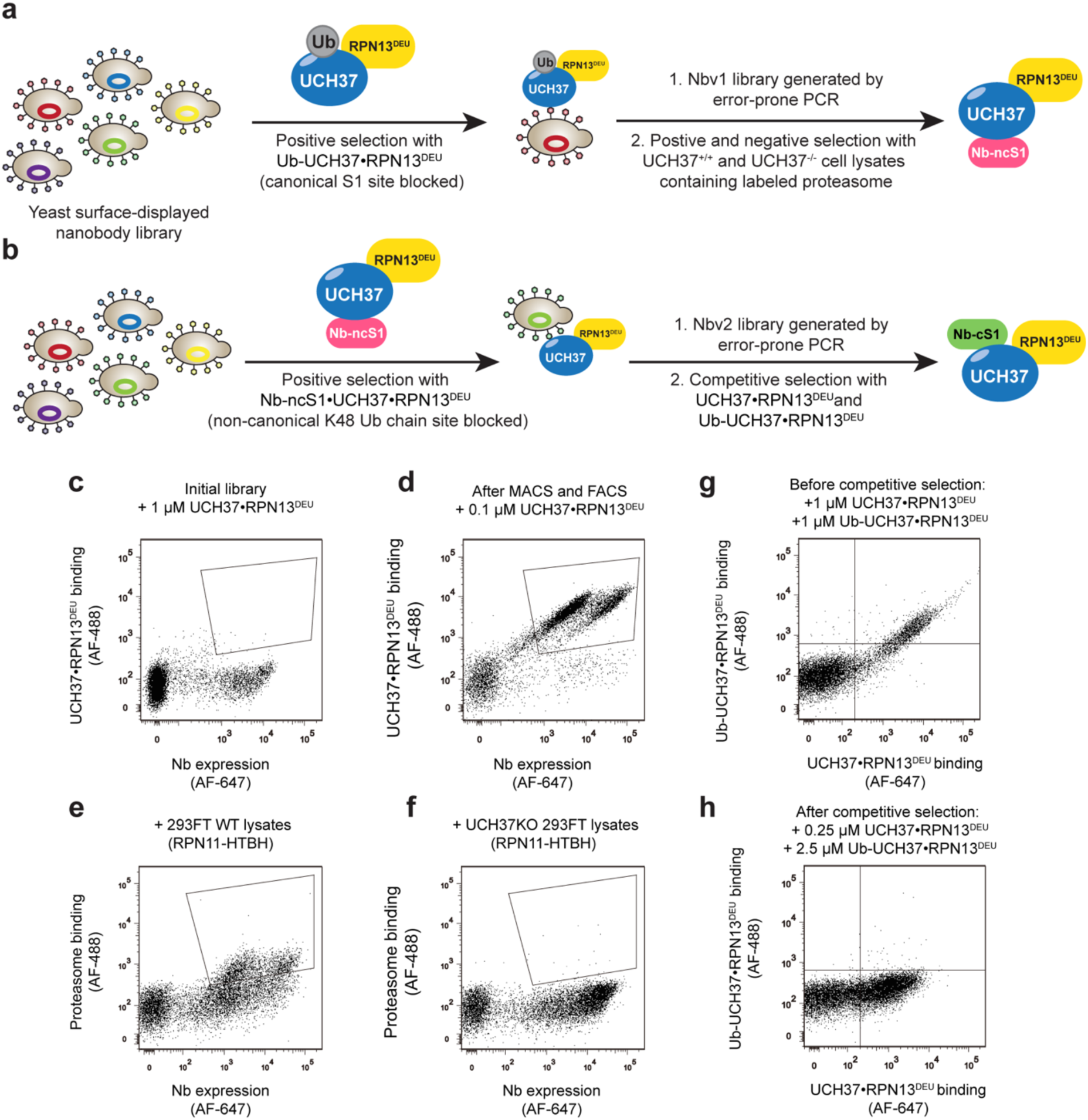
Screening strategy for the isolation of UCH37 nanobodies. (a) General scheme of Nb-ncS1 identification. Nb variants that bind to Ub-UCH37•RPN13^DEUBAD^ were enriched after several rounds of MACS and FACS, which include Nb-v1 as the most frequent clone. Nb-v1 was then diversified with error-prone PCR to generate a library for additional screening with HEK293FT or UCH37KO HEK293FT lysates containing biotinylated proteasomes. (b) General scheme of Nb-cS1 identification. Nb variants that bind to Nb-ncS1•UCH37•RPN13^DEUBAD^ were enriched after several rounds of MACS and FACS, which include Nbv2 as the most frequent clone. Nbv2 was then diversified with error-prone PCR to generate a library for additional screening with Flag-tagged UCH37•RPN13^DEUBAD^ (cS1 site free) and biotinylated Ub-UCH37•RPN13^DEUBAD^ (cS1 site blocked by Ub). (c-d) Scatter plots before (c) and after (d) MACS and FACS with biotinylated Ub-UCH37•RPN13^DEUBAD^ (d). Clones with high Nb expression and strong binding were enriched. (e-f) Scatter plots after positive and negative sorting against HEK293FT and UCH37KO HEK293FT lysates containing biotinylated proteasomes (RPN11-HTBH). Clones were enriched with high Nb expression and strong binding to WT proteasome (e) but not to UCH37KO proteasome (f). (g-h) Scatter plots before competitive sorting against Flag-tagged UCH37•RPN13^DEUBAD^ and biotinylated Ub-UCH37•RPN13^DEUBAD^. A large population of Nbs binds to both targets ((g), positive signal along the x-axis and y-axis). After competitive sorting, clones showed selective binding toward UCH37•RPN13^DEUBAD^ ((h), positive signal along x-axis, negative signal along y-axis).

After multiple rounds of sorting and sequencing, we identified an enriched clone, designated Nb-v1. Recombinant production and purification of Nb-v1 revealed a dissociation constant (*K*_d_) of 0.3 nM with free UCH37 (Supplementary Figure 2b). Pulldown assays confirmed that transfected Nb-v1 efficiently precipitated UCH37 from cell lysates but did not co-elute with other UCH37-interacting partners, such as proteasomal subunits (Supplementary Figure 2d). To further improve its binding affinity, we generated a library of Nb-v1 variants through error-prone PCR and conducted further rounds of selection targeting UCH37 within the context of the proteasome.

Next-generation sequencing (NGS) of the sorted population revealed several variants with enhanced binding to proteasomes from wild-type (WT) cells but not from UCH37-knockout cells (Figures 1e-f). Purification and binding analysis identified a variant with four substitutions (Q1R, L80Q, H99Y, A109P) that selectively binds UCH37 on 26S proteasomes (Supplementary Figure 2d, Supplementary Table 2), which we termed the non-canonical S1 (Nb-ncS1) binder (see below).

We then incubated the yeast-displayed nanobody library with the biotinylated Ub-UCH37•RPN13^DEUBAD^ complexed with Nb-ncS1 to block the non-canonical K48 binding site, thereby increasing the likelihood of isolating nanobodies that bind the canonical S1 site (Figure 1b). After enriching the population and identifying a high-frequency clone (Nb-v2), we performed affinity maturation using error-prone PCR, similar to the process for generating Nb-ncS1. To screen for canonical S1 (cS1) site binders, we used both biotinylated Ub-UCH37•RPN13^DEUBAD^ and Flag-tagged UCH37•RPN13^DEUBAD^ complexes (Figure 1b). Binders to the cS1 site were expected to interact only with Flag-tagged UCH37•RPN13^DEUBAD^, as the biotinylated complex had its cS1 site blocked by Ub (Figures 1g-h). This approach identified three consensus mutations (L47F, T50I, K99R) that produce a highly selective binder for the canonical S1 site, designated as Nb-cS1 (Supplementary Table 2).

### Nanobodies Bind UCH37 with High Affinity

Pull-down assays were conducted to confirm the site-specific binding of the selected nanobodies. HaloTag-fused Nb-cS1 and Nb-ncS1, each immobilized on a chloroalkane resin, successfully pulled down recombinantly expressed UCH37•RPN13^DEUBAD^ (Figure 2a). However, only Nb-ncS1 bound UCH37•RPN13^DEUBAD^ when the canonical S1 site was blocked by covalently attaching Ub to the active site cysteine (Figure 2a). These results indicate that the selection strategies were effective, with Nb-cS1 targeting the canonical Ub binding site and Nb-ncS1 engaging a distinct region outside the primary S1 site.

**Figure 2:**
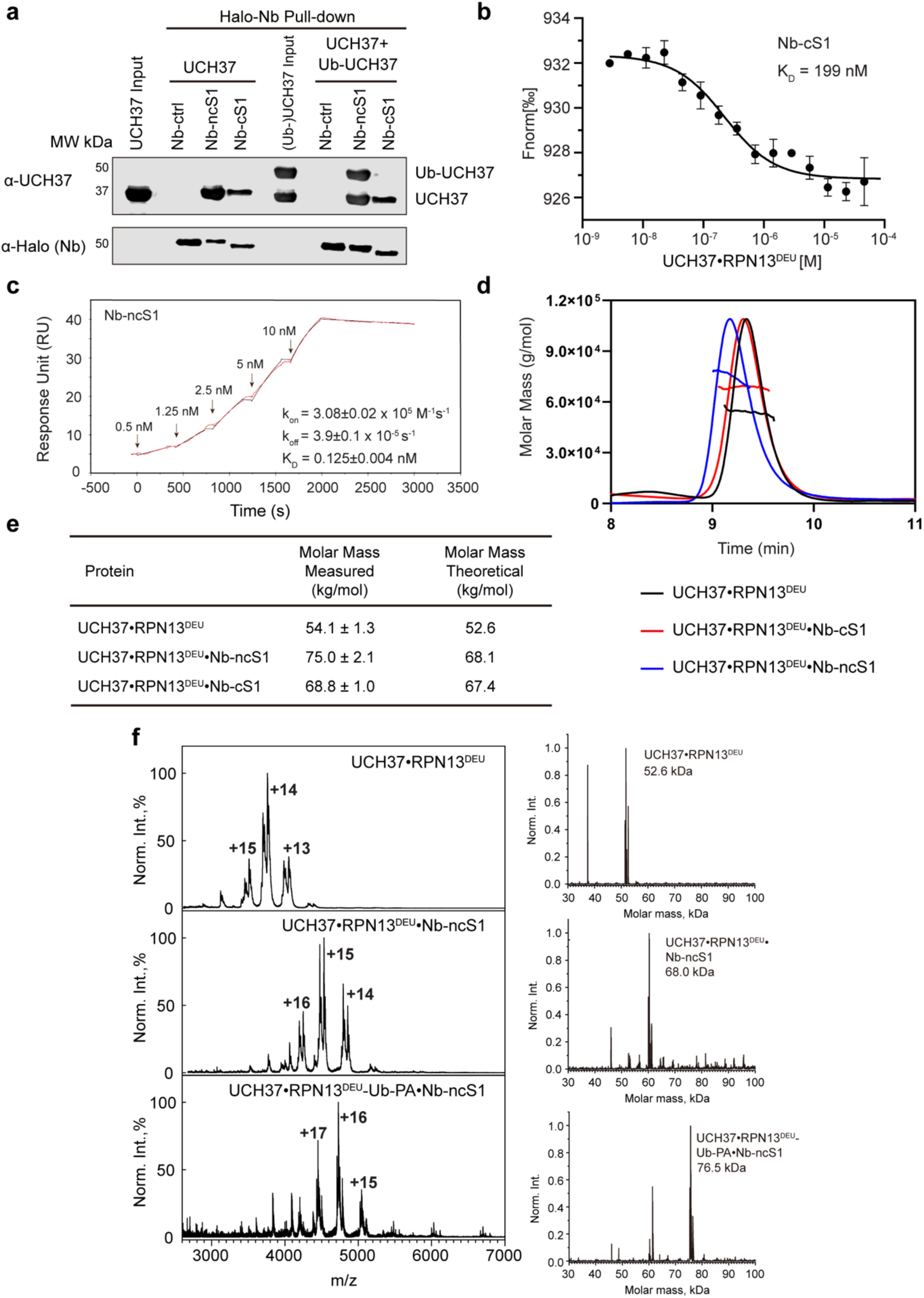
Nb-ncS1 and Nb-cS1 Bind to UCH37 with High Affinities. (a) Western blot analysis of Halo-tagged nanobody pull-downs with UCH37•RPN13^DEUBAD^ or a mixture of UCH37•RPN13^DEUBAD^ and Ub-UCH37•RPN13^DEUBAD^. Nb-ncS1 pulls down UCH37•RPN13^DEUBAD^ whether it’s conjugated to Ub or not, while Nb-cS1 can only pull down UCH37•RPN13^DEUBAD^. Immunoblotting was performed with the α-UCH37 antibody and α-Halo tag antibody. (b) Microscale thermophoresis (MST) experiments showing affinity between Nb-cS1 and UCH37•RPN13, n = 2 independent measurements, error bars represent the standard deviation. (c) Single-cycle surface plasmon resonance (SPR) experiment showing the interaction between Nb-ncS1 and UCH37. Experimental data is in red, and the curve fit is in black. (d) SEC-MALS elution profile of UCH37•RPN13^DEUBAD^(black) or UCH37•RPN13^DEUBAD^ incubated with Nb-ncS1(blue) or Nb-cS1(red). (e) Theoretical and measured molecular weight (kDa) of UCH37•RPN13^DEUBAD^ and its complex with Nb-ncS1 or Nb-cS1 obtained from SEC-MALS measurements. (f) Native ESI-MS spectra of free UCH37•RPN13^DEUBAD^ (top panel), Nb-ncS1-bound UCH37•RPN13^DEUBAD^ (middle panel), and Nb-ncS1 bound to Ub-UCH37•RPN13^DEUBAD^ (bottom panel). Additional peaks observed in the +13 to +15 charge states for free UCH37•RPN13^DEUBAD^ and in the +14 to +16 charge states for the complex correspond to N-/C-terminal truncations in the DEUBAD domain of RPN13. Deconvoluted spectra corresponding to each raw trace are shown to the right.

To quantify binding affinities, we used microscale thermophoresis (MST) and surface plasmon resonance (SPR). MST revealed that Nb-cS1 binds UCH37•RPN13^DEUBAD^ with a *K*_d_ of 0.2 μM (Figure 2b), while Nb-ncS1 exhibited a significantly higher affinity (*K*_d_ = 0.15 nM) (Supplementary Figure 2a). Although the concentrations of Nb-ncS1 and UCH37•RPN13^DEUBAD^ used in MST were outside the optimal range relative to the expected *K*_d_, SPR confirmed a similar affinity for Nb-ncS1 binding to UCH37 alone, with a *K*_d_ of 0.125 nM (Figure 2c).

We also assessed the binding stoichiometry of the nanobody complexes using size-exclusion chromatography coupled with multi-angle light scattering (SEC-MALS) and native electrospray ionization mass spectrometry (ESI-MS). Upon incubation with Nb-ncS1 or Nb-cS1, the UCH37•RPN13^DEUBAD^ complex exhibited shifts in both retention time and molar mass by SEC-MALS (Figure 2d). The measured molar masses of the complexes with Nb-ncS1 and Nb-cS1 were 75 kDa and 69 kDa, respectively (Figure 2e), consistent with a 1:1 binding stoichiometry for each nanobody. However, the ∼7 kDa discrepancy between the measured and theoretical mass for the Nb-ncS1 complex prompted further analysis by ESI-MS. Native mass spectrometry confirmed the formation of a well-defined 1:1 complex between UCH37•RPN13^DEUBAD^ and Nb-ncS1 (Figure 2f). Notably, native MS also showed that when the canonical S1 site was covalently occupied by Ub-PA, Nb-ncS1 still bound and formed a stable 1:1 complex with UCH37, consistent with its engagement at a distinct binding site.

### Nanobodies Bind Distinct Sites on UCH37

Having identified high-affinity nanobodies, we next sought structural insights into their binding sites. We successfully solved the X-ray crystal structure of UCH37·RPN13^DEUBAD^ in complex with Nb-cS1 at a resolution of 2.40 Å (Figure 3a, Supplementary Table 3). Superimposing this structure with the previously reported Ub-conjugated UCH37·RPN13^DEUBAD^ complex revealed that Nb-cS1 occupies the canonical S1 site, overlapping with the position of Ub (Figure 3b). In both complexes, UCH37 residue W36 plays a central role in binding: in the nanobody-bound structure, W36 is inserted into a hydrophobic pocket formed by Y37, I50, A96, I101, A104, and W106 of Nb-cS1 (Figure 3c), similar to how it is contacted by Ub in the previously reported structure^32^. Despite this shared interface, several conformational differences distinguish the Nb-cS1- and Ub-bound forms. Most notably, the C-terminal helical domain of UCH37 is tilted in the Nb-cS1-bound structure, and the loop containing residue I216—which contacts I36 of Ub—is flipped (Figure 3b).

**Figure 3:**
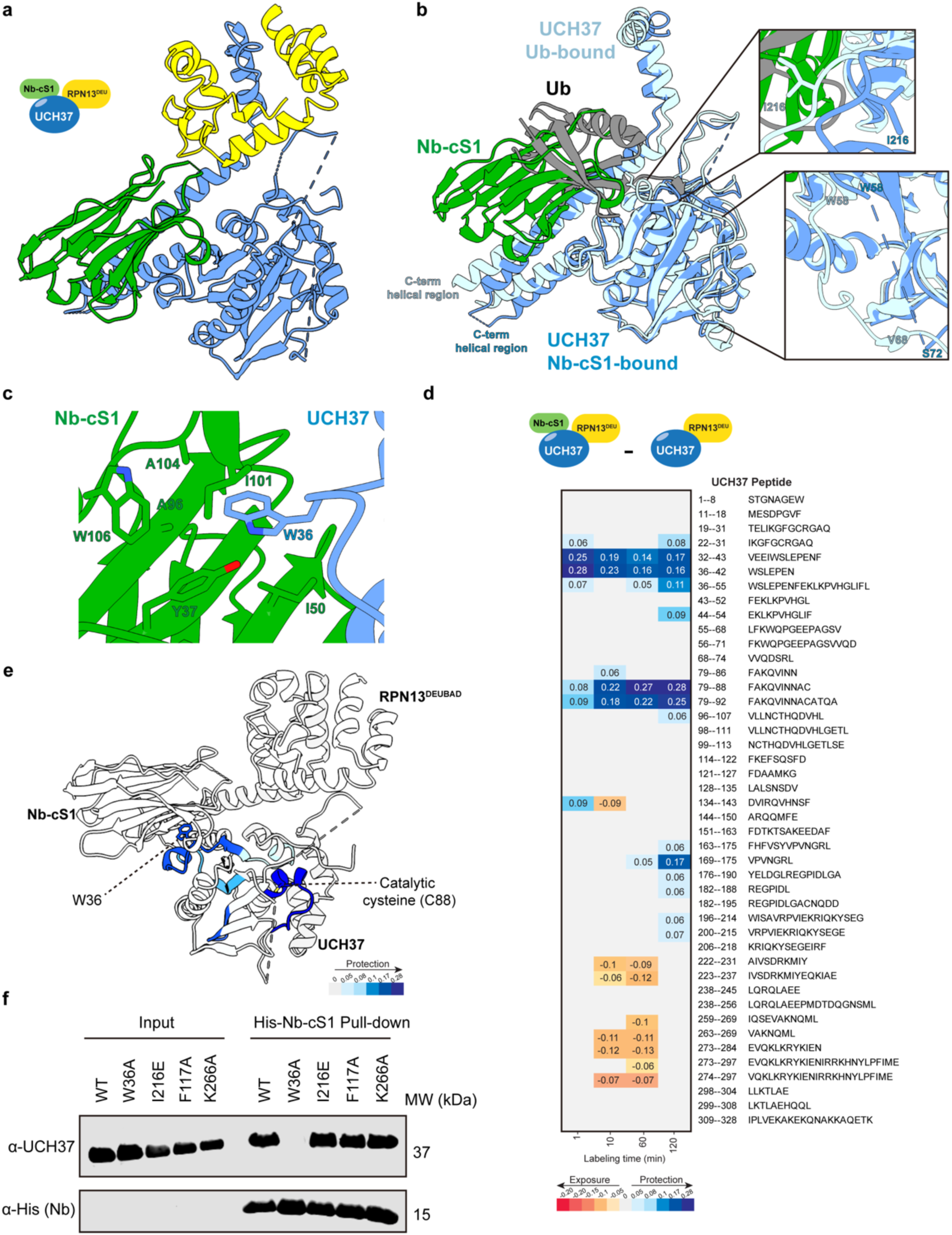
Structure of Nb-cS1 bound to UCH37•RPN13^DEUBAD^. (a) X-ray structure of Nb-cS1 (green) bound to UCH37 (blue) in complex with RPN13^DEUBAD^ (yellow). (b) Overlay of Nb-cS1-bound and Ub-bound (PDB: 4uel) structures with RPN13^DEUBAD^ omitted for clarity. Highlighted are the differences in the C-terminal helical regions of UCH37, as well as the loop containing I216 and residues 60–71. (c) Interaction of UCH37 W36 (blue) with a pocket in Nb-cS1 consisting of largely hydrophobic residues (green). (d) Differential deuterium uptake plot comparing UCH37•RPN13^DEUBAD^ in the presence and absence of Nb-cS1. Each data point represents the difference in deuterium uptake between the Nb-cS1-bound and free forms of the complex at the indicated time point. (e) Structure of UCH37•RPN13^DEUBAD^ bound to Nb-cS1 showing regions with statistically significant differences in deuterium exchange upon noncovalent binding of Nb-cS1. The data correspond to 2 h of deuterium labeling. (f) Western blot analysis of 6xHis-tagged Nb-cS1 pull-downs with UCH37•RPN13^DEUBAD^ variants. UCH37 W36A reduces binding to Nb-cS1. UCH37 I216E reduces binding of Ub to the canonical Ub-binding site^32^. UCH37 F117A reduces binding of K48-linked Ub chains to the noncanonical Ub-binding site^29^. UCH37 K266A is a substitution located in the C-terminal helical domain. Immunoblotting was performed with the α-UCH37 antibody and α-His tag antibody.

Additionally, the region encompassing residues 60–71 is poorly resolved in the Nb-cS1-bound structure (Figure 3b), suggesting increased flexibility. These conformational differences may have implications for how UCH37 accommodates and edits polyUb chains and raised the question of whether nanobody binding, like Ub, influences the enzyme’s catalytic architecture.

In the apo form of UCH37, the catalytic cysteine C88 is misaligned with respect to other key residues, such as H164 and D179 (Supplementary Figure 3). Ub binding to the canonical S1 site induces a conformational rearrangement that brings C88 into proximity with H164 and D179 promoting catalytic competence^33^. Strikingly, the Nb-cS1-bound structure reveals that the nanobody similarly aligns the catalytic residues (Supplementary Figure 3), suggesting it stabilizes a catalytically competent conformation even in the absence of Ub.

To determine whether these crystallographic observations are preserved in solution, we used hydrogen-deuterium exchange mass spectrometry (HDX-MS) to compare the apo and Nb-cS1-bound forms of UCH37. The HDX-MS data revealed two regions of UCH37 with significantly reduced deuterium uptake upon Nb-cS1 binding (Figures 3d-e). The first, residues 32–43, includes W36—a key residue involved in binding both Ub and Nb-cS1—highlighting the shared engagement of this anchoring point. Functional relevance was confirmed by pulldown assays: mutating W36 to alanine (W36A) abolished binding to His-tagged Nb-cS1, whereas mutations in other ubiquitin-interacting regions—including I216E, which reduces Ub affinity in the S1 canonical binding site; F117A, which disrupts the K48 chain-specific backside interface; and K266A, located in a region of the ULD domain that undergoes faster hydrogen-deuterium exchange (see below; Figure 3d)—had no effect on Nb-cS1 binding (Figure 3f). The second protected region, spanning residues 79–92, encompasses the catalytic cysteine C88, supporting the conclusion that Nb-cS1 stabilizes the active conformation of the catalytic site.

In addition to these protections, HDX-MS also revealed modest increases in deuterium uptake in other regions during the intermediate labeling times (Figure 3d). These include the final β-strand of the catalytic domain and the adjacent N-terminal segment of the C-terminal helical domain (residues 222–237), as well as a second region within the helical domain (residues 263–284). These increases suggest enhanced flexibility in parts of the C-terminal helical domain, consistent with the conformational changes observed in the crystal structure. Functionally, this conformational remodeling may influence how UCH37 interacts with polyUb chains and RPN13. Supporting the impact on RPN13, western blot analysis (see below; Figure 6b) shows reduced co-elution of RPN13 with Nb-cS1-bound UCH37, suggesting that nanobody binding could perturb the interaction between UCH37 and its regulatory partner.

Obtaining an atomic-level structure of Nb-ncS1 bound to UCH37 proved challenging due to crystallization difficulties. Nevertheless, HDX-MS provided valuable insights into the binding interface and associated conformational changes. The most pronounced reduction in deuterium exchange occurred in residues 114–135 (Figures 4a–b), corresponding to the K48-linked ubiquitin chain-specific binding site on the backside of the enzyme. Outside this region, only minimal protection was observed at other known ubiquitin-binding surfaces, reinforcing the specificity of Nb-ncS1 for the backside interface. In addition, regions spanning residues 19–31, 55–74, and 200–218 showed moderate-to-strong protection, suggesting conformational stabilization of domains adjacent to the binding site. Together, these findings support a model in which Nb-ncS1 selectively engages the K48 chain-specific interface on the backside of UCH37.

**Figure 4:**
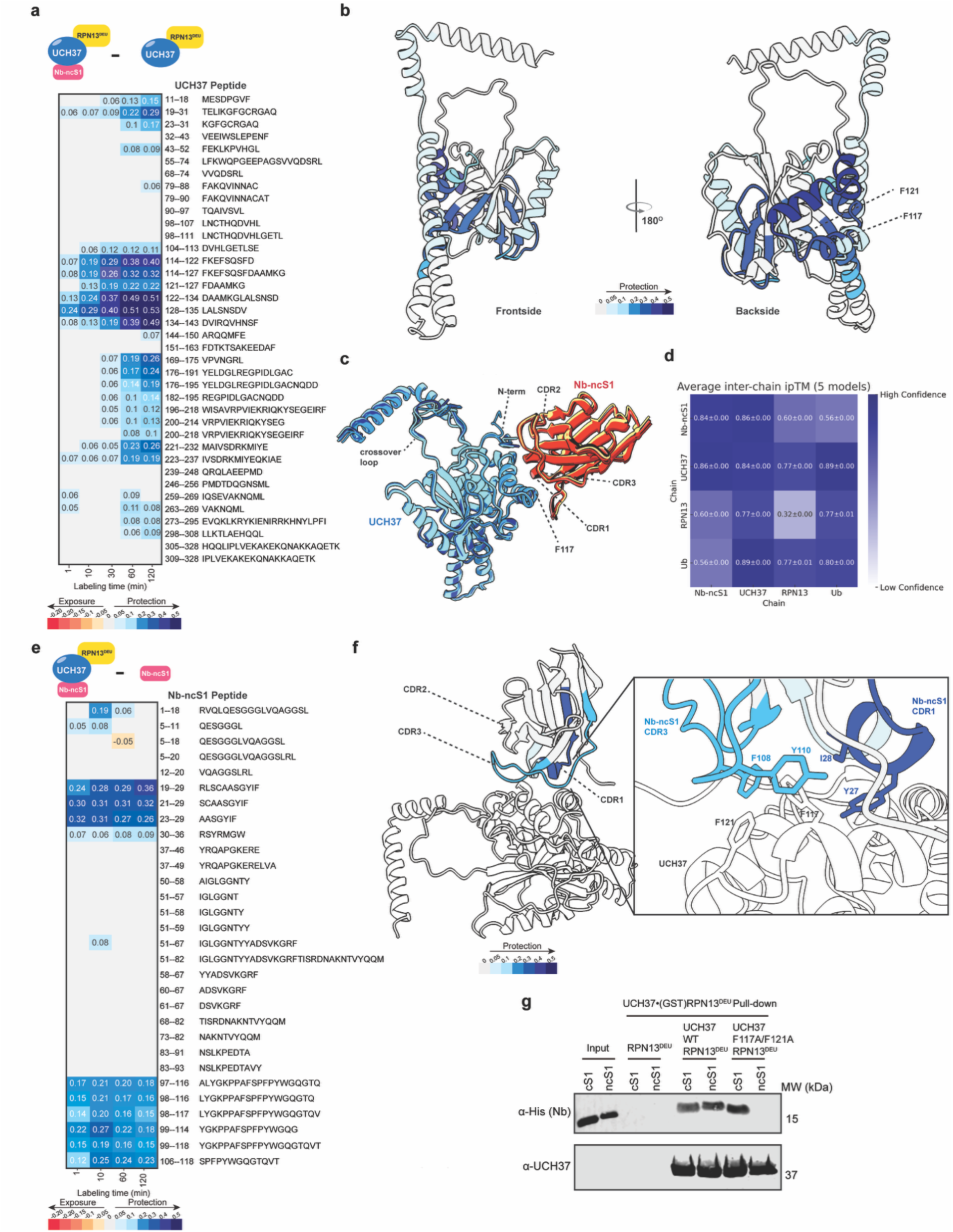
Nb-ncS1 is predicted to bind the aromatic-rich motif of UCH37. (a) Differential deuterium uptake plot comparing UCH37•RPN13^DEUBAD^ in the presence and absence of Nb-ncS1. Each data point represents the difference in deuterium uptake between the Nb-ncS1-bound and free forms of the complex at the indicated time point. (b) AlphaFold 3 model of UCH37 showing regions with statistically significant differences in deuterium exchange upon noncovalent binding of Nb-ncS1. The data correspond to 2 h of deuterium labeling. (c) Overlay of five AlphaFold 3 models of Nb-ncS1 bound to the Ub adduct of UCH37•RPN13. For clarity, Ub and RPN13 are omitted from the visualization. (d) Heat map showing the average inter-chain interface predicted TM-score (ipTM) across all five AlphaFold 3 models. The ipTM score for the Nb-ncS1 and UCH37 interface is 0.86, which indicates a high-confidence, high-quality prediction. (e) Differential deuterium uptake plot comparing Nb-ncS1 in the presence and absence of UCH37•RPN13^DEUBAD^. Each data point represents the difference in deuterium uptake between the UCH37•RPN13^DEUBAD^-bound and free forms of the complex at the indicated time point. (f) AlphaFold 3 model of Nb-ncS1 bound to UCH37•RPN13^DEUBAD^, highlighting regions with statistically significant changes in deuterium exchange upon noncovalent binding. The complementarity-determining regions (CDRs) 1 and 3 of Nb-ncS1 are predicted to engage UCH37 directly, forming hydrophobic contacts with the aromatic-rich motif on the backside of UCH37. (g) Western blot analysis of pulldowns with UCH37•(GST-tagged)RPN13^DEUBAD^ variants and either 6xHis-tagged Nb-cS1 or Nb-ncS1. UCH37 F117A/F121A reduces binding to Nb-ncS1, but not Nb-cS1. Immunoblotting was performed with the α-UCH37 antibody and α-His tag antibody.

AlphaFold 3 modeling further corroborated our experimental observations. To investigate the interaction between Nb-ncS1 and UCH37, we initially modeled full-length UCH37 in complex with RPN13 and Nb-ncS1. However, these initial models yielded low-confidence interface scores (interface predicted TM-score, ipTM ≈ 0.2) and incorrectly placed Nb-ncS1 at the canonical S1 ubiquitin-binding site—an outcome inconsistent with our yeast display selection strategy (Figure 1) and binding data (Figure 2), both of which indicate that Nb-ncS1 engages a distinct, non-canonical site. To refine the model, we incorporated ubiquitin into the canonical S1 site to sterically bias the prediction away from this region. Upon inclusion of ubiquitin, Nb-ncS1 consistently bound the backside of the enzyme across five independent models (Figure 4c), all exhibiting low predicted aligned error (PAE) (Supplementary Figure 4a) and high-confidence interface predictions. Notably, the ipTM scores for the Nb-ncS1–UCH37 interface exceeded 0.8 in all cases, indicating a robust and well-defined interaction in line with our HDX-MS data (Figure 4d). These models further predicted that complementarity-determining regions 1 and 3 (CDRs 1 and 3), but not CDR2, mediate engagement of the backside surface, specifically contacting the K48 chain-specific motif centered on F117 and F121 of UCH37 (Figure 4f).

To experimentally test whether CDRs 1 and 3, but not CDR2, contact UCH37, we again employed HDX-MS to compare the bound and unbound forms of Nb-ncS1. The data revealed two regions with markedly reduced deuterium uptake upon binding: residues 19-29, which comprises CDR1, and residues 99-120, encompassing CDR3 (Figures 4e-f). In contrast, no significant protection was observed for residues 52-69 which comprise CDR2. Thus, in agreement with the AlphaFold prediction, only CDRs 1 and 3 appear to mediate binding to UCH37.

Finally, to validate that Nb-ncS1 engages the predicted backside surface of UCH37, we tested the impact of mutating F117 and F121—two key aromatic residues within the K48-specific motif. Pull-down assays revealed a marked reduction in Nb-ncS1 binding to the F117A/F121A UCH37 mutant, while binding of Nb-cS1 remained unaffected (Figure 4g). Consistent with these findings, native ESI-MS detected no Nb-ncS1-bound form of the mutant UCH37 (Supplementary Figure 4b). Together, these results confirm that Nb-ncS1 selectively targets the non-canonical K48-specific site on the backside of UCH37.

### Nb-cS1 and Nb-ncS1 have Distinct Effects on UCH37

With the binding sites confirmed, we wanted to measure the effects on biochemical activities. Considering Nb-cS1 uses the canonical S1 Ub-binding site to engage UCH37, we surmised that it would interfere with the removal of small C-terminal Ub adducts while Nb-ncS1 would have little effect due to binding on the opposite face of the enzyme^32^. To test this, we examined the hydrolysis of Ub-aminomethylcoumarin (AMC) in the presence and absence of the nanobodies. As expected, the addition of Nb-cS1 significantly impairs Ub-AMC hydrolysis while Nb-ncS1 does not (Figure 5a). We also used Ub-propargylamine labeling as a secondary assay for validation. In this case, the C-terminal propargylamine adduct directly reacts with the active site cysteine of UCH37 forming a covalent Ub-UCH37 complex. Congruent with the results from the Ub-AMC assay, only Nb-cS1 exhibited significant inhibition (Figure 5b).

**Figure 5:**
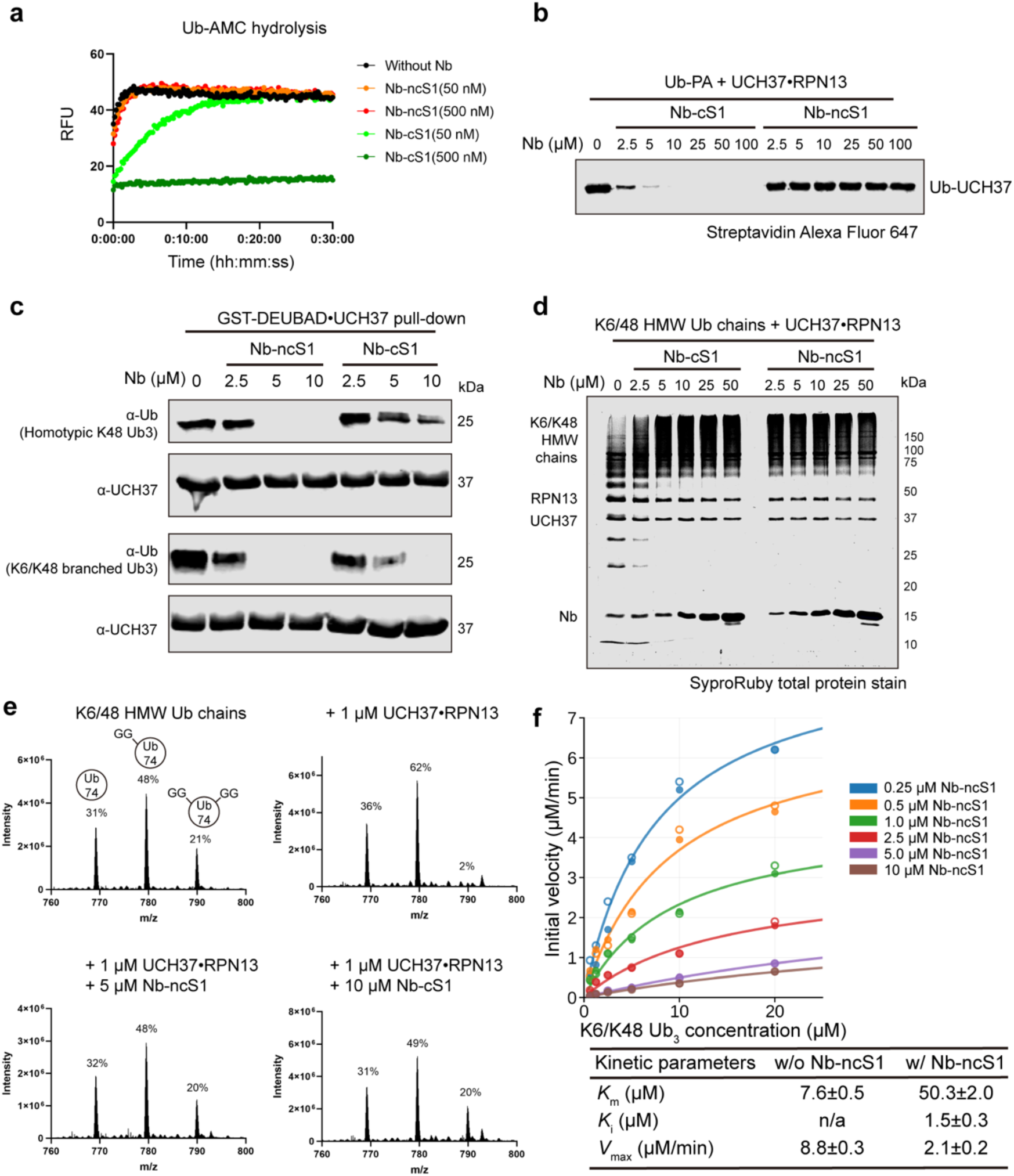
Nb-ncS1 and Nb-cS1 have different effects on the DUB activity of UCH37. (a) Hydrolysis of Ub-AMC (500 nM) by UCH37•RPN13^DEUBAD^ (2 nM) with the indicated concentrations of nanobodies. The released AMC fluorescence is monitored with excitation and emission wavelengths at 360 nm and 460 nm with a plate reader. (b) Western blot analysis of biotinylated Ub-propargylamine (2.5 μM) labeling by UCH37•RPN13^DEUBAD^ (2.5 μM) with the indicated concentrations of nanobodies. Immunoblotting was performed with Alexa Fluor 647-conjugated streptavidin. (c) Western blot analysis of UCH37•(GST-tagged)RPN13^DEUBAD^ pull-downs with homotypic K48 Ub3 or K6/K48 branched Ub3 (5 μM) with the indicated concentrations of nanobodies. (d) SDS-PAGE analysis of the cleavage of K6/K48 high-molecular weight (HMW) Ub chains (0.2 mg/ml) with UCH37•RPN13 (1 μM) co-incubated with different concentrations of nanobodies. Gels were imaged by SYPRO Ruby staining. (e) Middle-down MS analysis of HMW K6/K48 chains (0.2 mg/ml) subjected to UCH37•RPN13 (1 μM) co-incubated with different concentrations of nanobodies. Percentages correspond to the relative quantification values of the 11+ charge state for each Ub species: Ub1–74, 1xdiGly-Ub1–74, and 2xdiGly-Ub1–74. (f) Kinetic analyses of the cleavage of K6/K48 tri-Ub with UCH37•RPN13 (0.5 μM) with different concentrations of Nb-ncS1. Reactions were separated on a 15% SDS-PAGE gel and followed by SYPRO® Ruby staining. Initial velocities of Ub and di-Ub formation were converted to concentration (μM) per minute with Ub standards. These values were then fitted to the Michaelis-Menten equation using nonlinear regression in Prism. Nb-ncS1 exhibits mixed inhibition kinetics, with both apparent *K*m and *V*max changing with inhibitor concentration, indicating partial substrate access when the nanobody is bound. Error bars represent the standard deviation of two experiments.

Next, we sought to evaluate the impact of nanobodies on Ub chain binding. As we have previously shown, the non-canonical S1 (nc-S1) site located on the backside of UCH37 is necessary for binding K48-linked Ub chains^29^. Thus, not surprisingly, we found that Nb-ncS1 abolishes the binding of both unbranched and branched K48 chains in pull-down assays (Figure 5c). Nb-cS1 also diminished the binding of chains, but the effect was more pronounced for branched chains relative to unbranched chains. Work from our lab and others has demonstrated that the non-K48 linked Ub subunit of a branched chain forms contacts with UCH37^27,28^. It is likely these interactions overlap with the Nb-cS1 binding site, suggesting how the nanobody interferes with chain binding. Alternatively, as the crystal structure and HDX-MS data suggest, binding of Nb-cS1 to the canonical Ub-binding site could promote a conformation of UCH37 that might no longer be conducive to chain binding.

Since Nb-ncS1 and Nb-cS1 interfere with ubiquitin chain binding, we reasoned that each nanobody would also impede chain debranching. To test this, we added the nanobodies to reactions containing either model branched tri-ubiquitin substrates or enzymatically derived high–molecular weight (HMW) ubiquitin conjugates bearing branchpoints. Reactions were monitored by gel electrophoresis to detect the formation of lower–molecular weight ubiquitin conjugates, and by middle-down mass spectrometry to specifically assess the impact on branchpoints. Across these complementary approaches, we found that both Nb-ncS1 and Nb-cS1 inhibit chain debranching (Figures 5d–e, Supplementary Figure 5a–f).

More detailed quantitative analysis was performed for Nb-ncS1. Steady-state kinetic analysis of the debranching reaction using K6/K48-linked Ub_3_ revealed that Nb-ncS1 exhibits mixed inhibition, with both apparent *K*_m_ (5.2-fold increase) and *V*_max_ (4.7-fold decrease) changing significantly upon nanobody binding. The inhibition constant (*K*_i_ ≈ 1.5 μM) is ∼10,000-fold higher than the binding affinity (*K*_d_ ≈ 0.15 nM), indicating that substrate can still engage UCH37 when Nb-ncS1 is bound, albeit with reduced efficiency. This dramatic *K*_i_/*K*_d_ discrepancy is consistent with Nb-ncS1 binding to the ncS1 site on the face opposite the catalytic cleft, where it selectively disrupts the K48-specific substrate recognition surface while leaving other Ub-binding sites partially accessible. These findings support a model in which Nb-ncS1 interferes with optimal substrate positioning necessary for debranching without fully obstructing the catalytic core. In contrast, Nb-cS1 impairs both debranching and C-terminal hydrolase activity.

### Nb-ncS1 and Nb-cS1 Selectively Engage UCH37 in Cells

The use of Nb-ncS1 and Nb-cS1 as inhibitors of UCH37 requires that each nanobody selectively engage its target in cells. Before assessing proteome-wide selectivity, we first evaluated whether Nb-ncS1 and Nb-cS1 interact with UCH37 in a cellular context. HEK293FT wild-type (WT) and UCH37 knockout (UCH37KO) cells were transiently transfected with HA-Flag-tagged versions of Nb-ncS1, Nb-cS1, or a control nanobody (Nb-ctrl) (Figure 6a). Following immunoprecipitation, western blot analysis showed that Nb-ncS1 efficiently pulled down endogenous UCH37 from WT cells, along with proteasomal subunits including RPN13 (19S), RPN11 (19S), RPT2 (19S), and PSMA2 (20S) (Figure 6b). In contrast, while Nb-cS1 also co-immunoprecipitated UCH37, the amount recovered was substantially lower, consistent with a weaker binding interaction. Correspondingly, only trace amounts of RPN13 were detected, and no other proteasomal subunits were co-purified with Nb-cS1. No enrichment of UCH37 or other proteasomal components was observed from UCH37KO cells or when Nb-ctrl was used instead of Nb-ncS1 and Nb-cS1.

**Figure 6:**
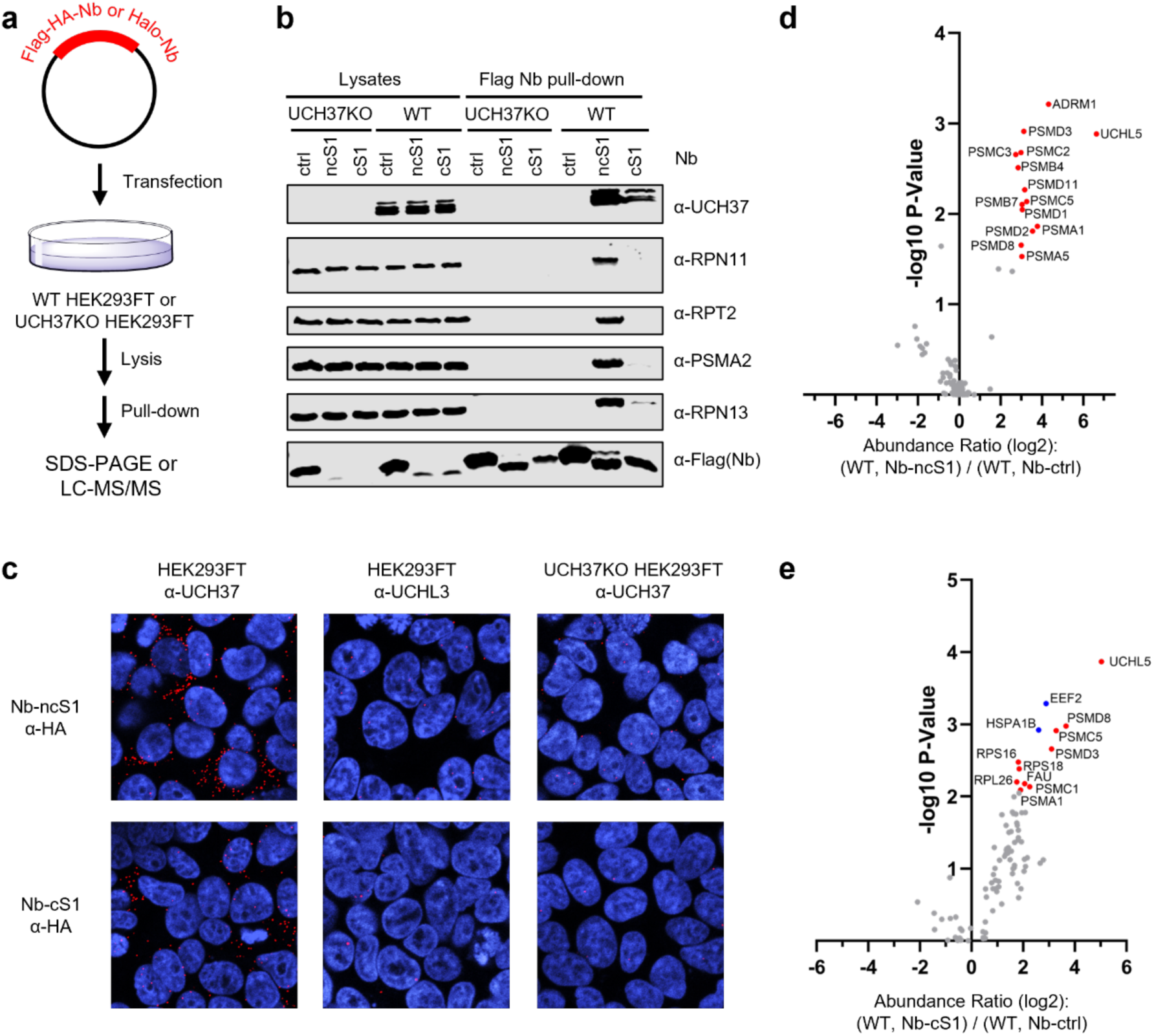
Nanobodies selectively interact with UCH37 in cells. (a) WT or UCH37KO HEK293FT cells were transfected with tagged nanobodies. Pulled-down proteins were analyzed by SDS-PAGE and LC-MS/MS. (b) Western blot analysis of anti-FLAG pull-downs with HEK293FT cell lysates transfected with FLAG tagged nanobodies. Samples were separated by SDS–PAGE and immunoblotted against UCH37 along with other proteasome subunits. (c) Proximity ligation assay (PLA) with HEK293FT cells transfected with different nanobodies. Cells were fixed and stained with the indicated antibodies followed by PLA. Blue: DAPI, RED: PLA signal. (d) Volcano plot from IP-TMT-MS data comparing proteins enriched by Halo-tagged Nb-ncS1 versus the Halo-tagged control nanobody (Nb-ctrl). UCH37 (UCHL5) along with other proteasomal subunits are shown in red. (e) Volcano plot from IP-TMT-MS data comparing proteins enriched by Halo-tagged Nb-cS1 versus the Halo-tagged control nanobody (Nb-ctrl). UCH37 (UCHL5) along with other proteasomal subunits and ribosomal proteins are shown in red.

To further probe cellular engagement, we employed a proximity ligation assay (PLA)^35^. In doing so, intracellular interactions between UCH37 and Nb-ncS1 or Nb-cS1 can be transformed into an amplifiable signal with oligonucleotide-labeled antibodies and rolling circle DNA replication. The signal is then detected by fluorescence microscopy using fluorescently-labeled oligonucleotides that bind to the amplified sequences. In HEK293FT cells expressing HA-Flag-tagged Nb-ncS1 or Nb-cS1, fluorescent puncta are observed upon the addition of antibodies that recognize UCH37 and the HA tag fused to the nanobodies (Figure 6c). By contrast, markedly fewer puncta are observed using the anti-HA antibody along with an antibody that recognizes UCHL3, another DUB in the same UCH sub-family as UCH37. Likewise, the expression of Nb-ctrl, which does not bind UCH37, also affords few puncta. Together with the pulldown data, these results demonstrate that Nb-ncS1 and Nb-cS1 interact with UCH37 in cells.

Having demonstrated cellular engagement, we sought to assess selectivity on a proteome-wide scale. The nanobodies were transiently expressed as Halo-tag fusions. Immunoprecipitation followed by mass spectrometry (IP-MS) analysis was then performed to identify interacting partners. Proteins were considered significantly enriched at p < 0.05 and log2 fold change > 1. In WT cells, the data revealed that UCHL5/UCH37 was indeed the most enriched target of Halo-Nb-ncS1 and Halo-Nb-cS1 relative to Halo-Nb-ctrl (Figures 6d-e). A significant number of 19S and 20S proteasome subunits were also identified, consistent with western blot analysis. Conversely, fewer proteasome subunits were detected in the Halo-Nb-cS1 immunoprecipitation (Figure 6e), while more ribosomal proteins were enriched—a pattern not observed in the Nb-ncS1 immunoprecipitation (Figure 6d). In UCH37KO cells, we did not detect enrichment of any proteasome components, indicating that UCH37 is required for both Halo-Nb-ncS1 and Halo-Nb-cS1 to associate with the proteasome (Supplementary Figure 6a–b).

Both nanobodies demonstrated selectivity toward UCH37 in HEK293FT cells, with Nb-ncS1 showing a cleaner proteomic profile and more efficient recovery of proteasomal complexes. The differences in proteasome recovery between the two nanobodies likely reflect their distinct binding modes and the structural consequences of engaging different sites on UCH37. Notably, neither nanobody enriched proteasomal components in UCH37KO cells, confirming that UCH37 is required for proteasome association. These results establish that both nanobodies selectively engage UCH37 in cellular contexts, with Nb-ncS1 providing superior performance for cellular studies requiring intact proteasomal complexes.

## DISCUSSION

Here, we report two nanobodies, Nb-cS1 and Nb-ncS1, that selectively engage distinct Ub-binding surfaces on UCH37 and enable the independent perturbation of its two enzymatic activities for the first time. Nb-cS1 binds the canonical S1 site and blocks both C-terminal Ub hydrolysis and K48-linked chain debranching, while Nb-ncS1 binds the non-canonical K48-specific (ncS1) site and selectively impairs only chain debranching. No existing small-molecule inhibitor can achieve this functional separation, making these nanobodies unique precision tools for dissecting UCH37’s dual roles.

Interestingly, Nb-ncS1, while tightly binding to UCH37, did not inhibit C-terminal Ub hydrolysis, highlighting the spatial separation of UCH37’s functional domains. This specificity could be crucial for therapeutic targeting, as selective inhibitors might spare certain UCH37 activities while still impairing its ability to regulate degradation pathways associated with Ub chain debranching. This is particularly relevant given the challenge of achieving selectivity when targeting deubiquitinases (DUBs), many of which share conserved catalytic motifs36. Our nanobody-based approach offers an avenue to design more precise inhibitors.

Importantly, kinetic analysis revealed that Nb-ncS1 exhibits mixed inhibition rather than simple competitive inhibition, with both apparent Km and Vmax changing upon nanobody binding. The large discrepancy between the inhibition constant (*K*_i_ ≈ 1.5 μM) and binding affinity (*K*_d_ ≈ 0.15 nM) indicates that substrate can still engage UCH37 when Nb-ncS1 is bound, consistent with binding at a site distinct from the active site. This mixed inhibition pattern provides direct mechanistic evidence that Nb-ncS1 disrupts optimal substrate positioning for debranching while allowing partial substrate access.

In cellular assays, Nb-ncS1 consistently demonstrated greater potency and selectivity for UCH37 than Nb-cS1. Quantitative IP-MS revealed that Nb-ncS1 enriches UCH37 more robustly than Nb-cS1, with minimal off-target binding, suggesting tighter and more specific engagement in cells. These findings are consistent with our biochemical analyses, which showed that Nb-ncS1 has higher affinity for UCH37 and selectively impairs chain debranching without broadly inhibiting other DUBs. In contrast, Nb-cS1 exhibited weaker cellular enrichment of UCH37 and modest off-target interactions, likely reflecting its lower binding affinity and broader inhibitory profile. Together, these data indicate that Nb-ncS1 is a more effective and selective tool for probing UCH37 function in the context of the proteasome and cellular proteostasis.

These tools open several important research directions. Structural studies of the INO80 complex reveal that the NFRKB recruitment subunit positions directly adjacent to UCH37, with its surface potentially occluding the same aromatic-rich ncS1 site (centered on F117 and F121) that Nb-ncS1 targets for K48-chain recognition. This spatial arrangement suggests that NFRKB binding may sterically prevent Nb-ncS1 access to its binding epitope when UCH37 is incorporated into the INO80 chromatin remodeling complex. If confirmed, this model would predict that Nb-ncS1 selectively inhibits only the proteasome-associated pool of UCH37, leaving the INO80-bound population functionally intact. Such selectivity would provide an unprecedented opportunity to dissect the distinct roles of UCH37’s debranching activity in proteasomal protein degradation versus chromatin remodeling contexts.

Testing this prediction could resolve long-standing questions about whether UCH37’s enzymatic activities contribute differently to its dual cellular functions, and whether therapeutic targeting of UCH37 could be achieved with context-specific precision rather than global inhibition.

In summary, Nb-cS1 and Nb-ncS1 are site-specific nanobody tools that, together, resolve UCH37’s two Ub-processing activities into independently addressable functions. The structural and biochemical basis for their selectivity, and the prediction that Nb-ncS1 acts only on the proteasome-bound pool of UCH37, position these nanobodies as precise probes for dissecting UCH37 biology at the proteasome and distinguishing it from UCH37’s role at INO80.

## MATERIALS AND METHODS

### Cell culture

Saccharomyces cerevisiae strain EBY100 (MATa AGA1::GAL1-AGA1::URA3 ura352 trp1 leu2delta200 his3delta200 pep4::HIS3 prbd1.6R can1 GAL) and yeast surface display vector, pCT used in yeast display were kindly provided by Dr. Sarah Moore (Smith College).

HEK293 stably expressing RPN11-His6-TEV-biotin tag-His6 (HEK293RPN11-HTBH) (Wang et al., 2007), UCH37KO HEK293RPN11-HTBH, and HEK293FT were cultured in DMEM medium (Genesee Scientific) supplemented with 10% fetal calf serum (Genesee Scientific), 100 units/ml penicillin and 100 μg/ml streptomycin at 37 °C in a humified atmosphere with 5% CO_2_.

### Plasmid and cloning

His8-MBP-TEV-UCH37 (Isoform 3) in the pVP16 vector was obtained from DNASU. His8-MBP-TEV-FLAG-UCH37 was cloned by inverse PCR with primers containing the FLAG tag. GST-3C-RPN13^DEUBAD^ (or RPN13 aa 253-407) was cloned from His10-RPN13 in pET19b, which was a gift from Joan Conaway & Ronald Conaway (Addgene plasmid # 19423). 1436 pcDNA Flag HA was a gift from William Sellers (Addgene plasmids #10792). Nanobodies were cloned into pET28b (Novagen) For bacterial expression, and 1436 pcDNA3 Flag HA vector for mammalian expression. For Halo-tagged nanobodies, the Halo tag was introduced at the N-terminus for Nb-ncS1, and C-terminus of Nb-cS1. All PCR amplifications were performed with Phusion® High-Fidelity DNA polymerase (New England Biolabs). The primers used in this study were purchased from IDT DNA.

### Construction of the synthetic nanobody (Nb) library

Our Nb library design is based on a published synthetic Nb library with slight modification and is displayed through the conventional aga1p-aga2p yeast surface display platform^30^. The Nb library was constructed by overlapping extension PCR of five oligonucleotides which composed the framework structure (oligonucleotides a and e) and randomized regions (oligonucleotides b, c, and d_short_/ d_medium_/ d_long_, and e) of Nb. The oligonucleotides were generated by performing PCR using a set of ten primers (a1_f + a2_r, b3_f + b4_r, c5_f + c6_r, d7_f + d8_short_r, or d8_medium_r, or d8_long_r, and e9_f+e10_r (Supplementary Table 1). The resulting mixture (oligonucleotides a, b, c, d, and e) was mixed at equal ratios for the following overlapping extension PCR; among them, oligonucleotides d were further composed of d_short_, d_medium_, and d_long_ mixed at a ratio of 1:1:2 to introduce different lengths of CDR3. Both PCRs were performed using Phusion® high fidelity DNA polymerase (New England Biolabs). The combined Nb gene library pool was further amplified using pCT_nb_PstI_f and pCT_nb_BamHI_r (Supplementary Table 1). 10 μg of the amplified Nb gene library was transformed to the yeast cells with 2 μg of the Nhel and BamHI (New England Biolabs) digested pCT plasmid through electroporation (Bio-Rad Gene Pulser Xcell). Dilutions of transformed yeast were then plated on a dropout medium without tryptophan (–Trp) as single colonies to obtain an estimate of library diversity. The diversity of the Nb library was further validated through next-generation sequencing (NGS).

### Construction of the random mutagenized Nb-v1 and Nb-v2 library

The **Nb-v1** and **Nb-v2** gene was randomly mutagenized using Mutazyme II error-prone DNA polymerase (Agilent Technologies). The mutagenized gene library pool was then amplified with the same primers using Phusion ® high-fidelity DNA polymerase (New England Biolabs). To prepare electrocompetent yeast cells, a 50 ml culture of EBY100 cells was grown to OD 1.5 and treated with 25 mM DTT for 15 mins. The cells were then harvested, washed, and resuspended in 2 x 150 μl of ice-cold 10 mM Tris, pH 7.5, 270 mM sucrose, 2 mM MgCl_2_. Cells were transformed separately using 2 μg of NheI and BamHI (New England Biolabs) of digested pCT vector and 10 μg of amplified Nb gene library pool through electroporation. The combined transformed cells were recovered in YPD media before being exchanged into SDCAA media. Recovered cells were serially diluted and plated in -Trp dropout media for 2-4 days at room temperature to estimate the size of the library.

### Next generation sequencing

The sequencing library was generated through reduced cycle amplification to add Illumina adapters to the Nb in two sequential PCR. The libraries were pooled with 40% Phi-X library spike-in (Illumina) and sequenced on Illumina MiSeq using Nano v2-500 cycle kit, with 251 bp paired-end chemistry. The quality of the sequence data was checked using FastQC. For data analysis, the paired-end reads were merged using FLASH. The resulting reads were trimmed by BBDuk and translated with SeqKit. Downstream analysis was performed in R. The enrichment ratios of specific variants were calculated by dividing the normalized Nb frequencies enriched through sorting over the normalized Nb frequencies presented in the initial error-prone Nb-ncS1 library.

### Protein purification

All protein purifications were performed at 4 °C. Biotin-Ubiquitin_1-75_-propargylamine (Biotin-Ub-PA) was synthesized from Avi-tagged Ub_1-75_-intein. Briefly, Avi-Ub1-75-intein was expressed in E. coli Rosetta (DE3) pLysS using 2xYT media. The cells were grown at 37 °C to OD 0.6 before being induced with 0.5 mM IPTG at 18 °C overnight. The cells were lysed in 20 mM sodium phosphate, pH 6.0, 500 mM NaCl, and 1 mM EDTA. The Avi-Ub_1-75_-intein was loaded onto chitin resin (New England Biolabs). The immobilized protein was washed and then eluted with lysis buffer in the presence of 100 mM sodium 2-mercaptoethanesulfonate (MESNA) (Acros Organic) for 48 hrs at 4 °C to generate Avi-Ub1-75-MESNA through on-resin cleavage. The Avi-Ub1-75-MESNA (final concentration was 500 μM) was incubated in a buffered solution containing 20 mM Tris, pH 8.0, 10 mM N-hydroxysuccinimide, 5 mM propargylamine (Alfa Aesar) at 37°C for overnight to yield AviUb-PA.

Complexes of UCH37 and RPN13 or RPN13^DEUBAD^ were expressed and purified as previously described^27^, but with a slight modification. Briefly, His-MBP-TEV-UCH37 and His-RPN13 or GST-3C-RPN13^DEUBAD^ constructs were expressed in BL21(DE3) pLysS E. coli cells grown at 37°C in LB media. The cells were grown at 37 °C to OD 0.8 before being induced with 0.4 mM IPTG at 20 °C. Cultures were harvested after 16 hr and frozen at – 80 °C. UCH37 and RPN13 or RPN13^DEUBAD^ cell pellets are mixed at 1:1 prior to lysis. Cell pellets were resuspended in lysis buffer (50 mM HEPES pH 7.4, 150 mM NaCl, 1 mM EDTA, and 1 mM TCEP), lysed by sonication, and clarified by centrifugation. UCH37•RPN13 and UCH37•RPN13^DEUBAD^ were purified following different chromatographic steps.

For UCH37•RPN13, clarified lysate was incubated with amylose resin for 2 hr at 4 °C, washed with lysis buffer, eluted with lysis buffer containing 10 mM maltose, but without EDTA, and then incubated overnight with TEV protease at 4°C to remove His-MBP tag. Then TEV protease-cleaved product was incubated with Ni-NTA resin for 1 hr, washed with Ni-NTA lysis and wash buffer (50 mM HEPES pH 7.4, 150 mM NaCl, 10 mM imidazole, and 0.5 mM TCEP), and eluted with the Ni-NTA elution buffer (50 mM HEPES pH 7.4, 150 mM NaCl, 300 mM imidazole, and 0.5 mM TCEP).

For UCH37•RPN13^DEUBAD^, clarified lysate was incubated with Ni-NTA resin for 2 hr at 4°C, washed with Ni-NTA wash buffer, and eluted into Ni-NTA elution buffer, and then incubated overnight with TEV protease. The TEV protease cleaved product was then incubated with GST resin for 1 hr, washed with lysis buffer (50 mM HEPES pH 7.4, 150 mM NaCl, and 1 mM TCEP), and eluted with GST elution buffer (GST lysis buffer plus 10 mM glutathione). Eluate was incubated overnight with 3C protease at 4°C. For both UCH37•RPN13 and UCH37•RPN13^DEUBAD^, the eluate was concentrated and separated on a Superdex 200 (GE) gel filtration column.

The His-Halo-nanobody/His-nanobody (Nb) variants were purified through immobilized metal affinity chromatography. The His-tagged Nbs were expressed in E. coli BL21 (DE3). The cells were grown at 37°C to OD ∼0.6 before being induced at 20°C with 0.4 mM IPTG overnight. The harvested cells were lysed in 50 mM HEPES, pH 7.5, 300 mM NaCl, 10 mM imidazole, and 0.1% Triton X-100 and purified using Ni-NTA (GoldBio) affinity chromatography. The elution buffer was 50 mM HEPES, pH 7.5, 300 mM NaCl, 300 mM imidazole. The Nbs were further purified through size exclusion chromatography using a HiLoad Superdex 75 pg on an ÄKTA FPLC (GE Healthcare). The running buffer was 50 mM HEPES, pH 7.5, 300 mM NaCl, 0.5 mM TCEP.

E1, UBE2D3, UBE2R1, UBE2N/UBE2V2, UBC1, and UBE3C were purified as previously described with slight modification^27,39–42^. Briefly, E1, UBE2D3, UBE2R1, and UBE2N/UBE2V2 constructs were expressed in Rosetta 2(DE3)pLysS Escherichia coli cells grown at 37°C in LB media. Once cells reached an OD600 of 0.6–0.8, 400 μM Isopropyl ß-D-1-thiogalactopyranoside (IPTG) was added, and the temperature was reduced to 20°C to induce protein expression. Cells were then harvested after 16 hr and resuspended in Ni-NTA lysis buffer (50 mM Tris pH 7.5, 300 mM NaCl, 1 mM EDTA and 10 mM imidazole), lysed by sonication, and clarified by centrifugation. Clarified lysate was incubated with Ni-NTA resin for 2 hr at 4°C, washed with Ni-NTA lysis buffer, and eluted into Ni-NTA elution buffer (Ni-NTA lysis buffer plus 300 mM imidazole).

NleL (aa 170–782) was purified as previously described^27,43^, but with a slight modification. Briefly, NleL was expressed in BL21(DE3)pLysS E. coli cells grown at 37°C in LB media. Once cells reached an OD600 of 0.6, IPTG (200 μM) was added, and the temperature was reduced to 20°C to induce protein expression. Cells were then harvested after 16 hr and resuspended in GST lysis buffer (50 mM Tris pH 8.0, 200 mM NaCl, 1 mM EDTA, and 1 mM DTT), lysed by sonication, and clarified by centrifugation. Clarified lysate was then incubated with GST resin for 2 hr at 4°C, washed with GST lysis buffer, and eluted into GST elution buffer (GST lysis buffer plus 10 mM reduced glutathione). Eluate was concentrated in TEV protease buffer (50 mM Tris pH 8.0, and 0.5 mM TCEP), cleaved overnight with TEV protease, and further purified using anion exchange chromatography.

### Generation of biotinylated Ub conjugated UCH37•RPN13^DEUBAD^

To prepare biotinylated Ub conjugated to UCH37•RPN13^DEUBAD^, the Avi-Ub-PA was biotinylated through BirA reaction before conjugated to UCH37•RPN13^DEUBAD^. Briefly, the Avi-Ub-PA (final concentration was 40 μM) was added to a solution containing 10 mM Tris, pH 8.0, 5 mM MgCl_2_, 2 mM ATP, 150 μM D(+) biotin (GoldBio), and 0.4 μM BirA at 37°C until the biotinylation reaction went to completion. The reaction was verified using MALDI-TOF MS. The 5 μM biotinylated Ub-PA was incubated with 2.5 μM UCH37•RPN13DEU in a buffered solution containing 50 mM HEPES, pH 7.5, 50 mM NaCl, 1 mM TCEP at room temperature for 1 hr with gentle rocking. The biotinylated ternary complex was further purified through size exclusion chromatography using a HiLoad Superdex 200 pg column.

### Synthesis of defined Ub chains

Native homotypic K48 chains were synthesized as previously described^27^. Briefly, 1 mM Ub, 300 nM E1, and 5 μM UBE2R1 were mixed in Ub chain synthesis reaction buffer (10 mM ATP, 10 mM MgCl_2_, 40 mM Tris-HCl pH 7.5, 50 mM NaCl, and 0.6 mM DTT) overnight at 37°C. Native K6/48 branched trimer was synthesized by mixing 2 mM K6/48R Ub, 1 mM UbD77, 300 nM E1, 5 μM UBE2D3, and 1 μM NleL in ubiquitin chain synthesis reaction buffer overnight at 37°C. Native K48/K63 branched trimer was made by first synthesizing K48 dimer by mixing 1 mM K48/K63R Ub, 1 mM UbD77, 300 nM E1, 5 μM UBE2R1, then the purified K48 dimer is mixed with 1 mM K48/K63R Ub, 300 nM E1, 3 μM UBE2N/UBE2V2 to synthesize K48/K63 branched trimer. All reactions for native chains were quenched by lowering the pH with the addition of ammonium acetate solution (pH 4.4). Enzymes were then precipitated through multiple freeze-thaw cycles and further purified using cation exchange chromatography.

### Generation of high molecular weight Ub chains

Native K6/K48 and K48/K63 HMW Ub chains were synthesized as previously described^27^. Briefly, K6/K48 Ub chains were synthesized by mixing 1 mM Ub, 200 nM E1, 5 μM UBE2D3, and 1 μM NleL in Ub chain synthesis reaction buffer and K48/K63 Ub chains were synthesized using 1 mM Ub, 150 nM E1, 10 μM UBE2R1, and 5 μM UBE2N/UBE2V2 in Ub chain synthesis reaction buffer. Reaction mixtures were all incubated at 37°C overnight. All Ub chains were purified using size exclusion chromatography (Superdex 75pg) to isolate HMW chains > 75 kDa.

### Selection of nanobodies against UCH37

The display of the Nb on the yeast surface was induced by resuspending 10x diversity of the Nb library in the SGCAA media at 20 °C. The induced cells were resuspended in PBSF buffer (0.1% (v/v) BSA in PBS buffer, pH 7.4) and subjected to two subsequent magnetic activated cell sorting (MACS) and two fluorescent activated cell sorting (FACS) to select Nbs to bind to Ub-UCH37•RPN13^DEUBAD^ (for Nb-ncS1 selection) or Nb-ncS1•UCH37•RPN13^DEUBAD^ (for Nb-ncS1 selection).

For MACS, the induced cells were negatively sorted against DynabeadsTM M-280 Streptavidin (ThermoFisher Scientific) before positively sorted against Dynabeads coated with biotinylated antigens in a tube setting with magnets. All MACS were performed at 4 °C for 2 hrs. For FACS, the induced cells were labeled with chicken anti-c-Myc (Exalpha biologicals INC) at 1:250 dilution for 1 hr at room temperature. The labeled cells were then incubated with varied concentrations of biotinylated antigens for 1 hr at room temperature before being simultaneously labeled with goat anti-chicken IgY (H+L) secondary antibody, Alexa FluorTM 647 and streptavidin Alexa Fluor 488 conjugate (ThermoFisher Scientific) at 1:100 dilution for 30 mins on ice in dark. Sortings were performed on a BD FACS ARIA II with 488 nm (530/30 band pass collection filter) and 640 nm (670/30 band pass filter) excitation lasers. At the end of FACS, yeast display plasmids were isolated from the enriched clones using the Zymoprep yeast plasmid miniprep kit (Zymo Research). The enriched clones were identified through random Sanger sequencing, which identified Nb-v1 (Nb-ncS1) or Nbv2 (Nb-cS1) as the most enriched clones.

### Selection of nanobodies against proteasome-associated UCH37(Nb-ncS1)

The Nb-v1 library generated by error-prone PCR was cultured, induced, and washed as described above. The induced yeast cells were first labeled with chicken anti-c-Myc at 1:250 dilution for 30 mins at room temperature. This was followed by the incubation of the cells with clarified cell lysate of the HEK293RPN11-HTBH in a buffered solution containing 50 mM HEPES, pH 7.5, 50 mM NaCl, 5 mM MgCl_2_, 2 mM ATP, 10% glycerol for 45 minutes at room temperature. The cells were then similarly labeled with the secondary antibodies and sorted through FACS as mentioned above. For negative sorting, the cell lysate of UCH37 KO HEK293TRPN11-HTBH was used as the antigen. The plasmids containing the gene of the enriched clones were extracted through the Zymoprep yeast plasmid miniprep kit and submitted for Sanger sequencing or used in the preparation of the NGS sequencing library.

### Competitive sorting of nanobodies against UCH37•RPN13^DEUBAD^ and Ub-UCH37•RPN13^DEUBAD^ (Nb-cS1)

The Nbv2 library generated by error-prone PCR was cultured, induced, and washed as described above. The induced yeast cells were incubated with both FLAG-tagged UCH37•RPN13^DEUBAD^ and biotinylated Ub-UCH37•RPN13^DEUBAD^ for 1 hr at room temperature. After washing, the cells are then incubated with rabbit anti-FLAG antibody (Cell Signaling Technology) for 30 mins at room temperature. This was followed by the incubation with Alexa Fluor 647-conjugated AffiniPure Goat Anti-Rabbit IgG (Jackson ImmunoResearch) and streptavidin Alexa Fluor 488 conjugate for 30 mins on ice in the dark. The cells were then sorted through FACS selecting positive 647 and negative 488 signal clones.

### In vitro pull-down assay

Halo-tagged nanobodies were immobilized on the pre-equilibrated HaloLink (Promega) resin for 2 hours at 4 °C with gentle rocking. After washing, the resin with immobilized nanobodies was incubated with the recombinantly expressed UCH37 RPN13^DEUBAD^, or also with Ub-UCH37•RPN13^DEUBAD^ for 2 hr at 4 °C with gentle rocking. The enriched proteins were extensively washed and eluted using 6x Laemmli loading buffer at 95°C for 3 mins before being subjected to SDS-PAGE and Western Blot analysis.

His-tagged nanobodies were immobilized on the pre-equilibrated Nickel-NTA resin (Genscript) for 2 hours at 4 °C with gentle rocking. After washing, the resin with immobilized nanobodies was incubated with the recombinantly expressed UCH37 WT or mutants in complex with RPN13^DEUBAD^ for 2 hours at 4 °C with gentle rocking. The enriched proteins were extensively washed and eluted using 6x Laemmli loading buffer at 95°C for 3 mins before being subjected to SDS-PAGE and Western Blot analysis. GST-tagged RPN13^DEUBAD^ in complex with UCH37 WT or UCH37 F117A/F121A were immobilized on the pre-equilibrated glutathione resin(Genscript) for 2 hours at 4 °C with gentle rocking. After washing, the resin with immobilized UCH37•RPN13^DEUBAD^ was incubated with 5 μM Nb-ncS1 or Nb-cS1. 5 uM homotypic K48 Ub3 or K6/48 branched Ub3 were also added in the ubiquitin chain competition pull-down. The enriched proteins were extensively washed and eluted using 6x Laemmli loading buffer at 95°C for 3 mins before being subjected to SDS-PAGE and Western Blot analysis.

Western blot analysis was imaged with Odyssey CLx Imager (LI-COR Biosciences) and performed using the following antibodies: rabbit anti-UCH37 (abcam, EPR4896), rabbit anti-Flag (Cell Signaling Technology, 14793S), rabbit anti-His tag (Cell Signaling Technology, 2365S). IRDye® 680RD goat anti-rabbit and IRDye® 800CW goat anti-mouse (LI-COR Biosciences) were used as the secondary antibodies.

### Microscale thermophoresis (MST)

MST was performed to study Nb-ncS1 or Nb-cS1 with UCH37•RPN13 interaction using Monolith NT.115 Microscale Thermophoresis instrument (Nano Temper Technologies, Germany). In brief, Nb-ncS1 or Nb-cS1 was labeled with Fluorescein-isothiocyanate (Sigma) according to the manufacturer’s instructions. Labeling reagents were removed by desalting column and buffer was exchanged into 50 mM Tris pH 7.4, 150 mM NaCl, and 10 mM MgCl_2_ with 0.05% tween-20. UCH37•RPN13 were serially diluted in the same buffer, to obtain varying concentrations from 4 µM to 0.12 nM (for Nb-ncS1) or 46 µM to 2.8 nM (for Nb-cS1) and Nbs were added with a final concentration of 20 nM (Nb-ncS1) or 50 nM (Nb-cS1). The binding study was carried out using NT.115 premium capillaries, and the assay was performed in duplicates. The data for MST were recorded at room temperature at 40% LED power and IR-laser power at 40%. Data analysis was performed using MO analysis software (Nano Temper). The Kd fit model is based on a molecular interaction with a 1:1 stoichiometry according to the law of mass action.

### Surface plasmon resonance (SPR)

The surface plasmon resonance experiments were performed using a BIACORE T200 (GE Healthcare) equipped with a Series S Sensor Chip NTA. The surfaces of the flow cells were conditioned using 350 mM EDTA for 1 min at a flow rate of 10 μl/min before injecting 0.5 mM NiSO_4_ for 1 min at 10 μl/min. The purified His-Nb (15 kDa) was then injected for 2 mins at 10 μl/min and immobilized on the Ni-NTA surface at a density of approximately 30 response units (RU). Single cycle kinetic titration experiments were performed by injecting purified UCH37 at increasing concentrations in 50 mM HEPES, pH 7.5, 50 mM NaCl, 50 μM EDTA, 0.005% (w/v) TWEEN 20 at a flow rate of 30 μl/min with a 300 s contact time and 900 s dissociation time. The flow cell surfaces were regenerated using 2 pulses of 350 mM EDTA for 2 min at a flow rate of 10 μl/min. The sensorgrams were double-referenced before analyzed using BIAevaluation (GE Healthcare). The association and dissociation rate constant, ka and kd, were determined from the direct curve fitting to a 1:1 binding model; the dissociation equilibrium constant, K_D_, was calculated from k_off_/k_on_. All experiments were performed at 25 °C.

### Size exclusion chromatograph-multi angle light scattering (SEC-MALS) analysis

SEC-MALS was performed in a TSKgel G3000SWXL column (TOSOH Biosciences) on a Agilent 1260 Infinity HPLC system (Agilent Technologies) coupled to a MALS detector, DAWN HELEOS II (Wyatt Technology) and a refractive index detector, Optilab T-rEX (Wyatt Technology). Nanobodies were incubated with UCH37•RPN13^DEUBAD^ on ice for 30 mins before the analysis. The flow rate was 1.0 ml/min and the mobile phase was 50 mM sodium phosphate, pH 6.5, 150 mM sodium chloride, and 10% isopropanol. Data was analyzed using Astra (Wyatt Technology) and the molar masses of the complexes were determined by the Zimm fit model. All the experiments were performed at room temperature.

### HDX-MS analysis

*Sample preparation.* To assess the deuterium uptake difference between free UCH37•RPN13^DEUBAD^ and Nb bound UCH37•RPN13^DEUBAD^, UCH37•RPN13^DEUBAD^ (45 μM) were pre-incubated with the Nb (90 μM) for 1 hr on ice in dilution buffer (50 mM HEPES, pH 7.4, 50 mM NaCl, and 0.5 mM TCEP). To assess the uptake differences between free Nb and UCH37•RPN13^DEUBAD^ bound Nb, UCH37•RPN13^DEUBAD^ (45 uM) were pre-incubated with Nb (45 uM) to avoid the presence of unbound Nb. All further steps, including deuterium labeling and reaction quench were performed on Trajan’s LEAP HDX platform (Trajan Scientific and Medical, Ringwood, Victoria, Australia). 4 μL of protein complex was diluted into either 96 μL of H_2_O buffer (20 mM sodium phosphate, pH 7.0 in H_2_O) for reference mapping or 96 μL of D_2_O buffer (20 mM sodium phosphate, pD 7.0 in D_2_O, 99.9% D_2_O from Cambridge Isotopes) to initiate deuterium labeling. After different times of exposure to the deuterated buffer at 15°C, 50 μL labeling reaction was quenched by mixing with 50 μL quenching buffer (0.8 M Guanidine-HCl, 1% acetic acid, pH 2.2) at 4°C. 50 uL of quenched samples were immediately injected into the LC-MS platform.

### HDX-MS data acquisition

The LC-MS data acquisition was performed on the ACQUITY UPLC M-Class System (Waters Corp.) and Synapt G2Si HDMS (Waters Corp.). Before the experiment, the MS instrument was calibrated in the positive ion mode using sodium iodide. The data was acquired in the positive mode in the mass range 100-2000 m/z using ESI with the following parameters: capillary voltage 2.5 kV, sampling cone 40 V, source offset 30 V, source temperature 80°C, desolvation temperature 175°C, cone gas 90 L/h, desolvation gas flow 600 L/h, nebulizer pressure 5 bar. To minimize hydrogen gas phase back exchange and keep the optimal separation of peptides within ion mobility cell, ion optics parameters were adjusted (StepWave transfer offset 15 V, DA1 3 V, DA2 0 V; IMS wave velocity 700 m/s, IMS wave height 40 V). Upon injection, samples were passed through an immobilized pepsin column (Enzymate BEH Pepsin Column, 2.1 × 30 mm, Waters Corp.) at 100 μL/min for inline digestion at 10°C. Following digestion, the resulting peptides were desalted on a ACQUITY UPLC HSS T3 VanGuard Pre-column (C18, 1.8 μm, 100 A, 2.1 × 5 mm, Waters Corp.) and separated on a ACQUITY UPLC HSS T3 microbore column (C18, 1.8 μm, 100 A, 1.0 × 50 mm, Waters Corp.) at 0°C with linear gradient of 80% B (A is 0.1% formic acid [FA] in H_2_O, and B is 0.1% FA in acetonitrile) over 12 min. The identification of peptides was performed using the data from the undeuterated sample acquired in MS^e^ mode, while HDX data was acquired using HDMS in resolution mode.

### HDX data analysis

Peptide identification was conducted using Protein Lynx Global Server (PLGS) version 3.0.1 software (Waters Corp). Data acquired from undeuterated samples were searched against the sequence database of proteins present in the complexes. Processing parameters include 556.2771 Da/e as lock mass for charge +1, lock mass window of 0.25 Da, and non-specific primary digest reagent. Data from peptides generated from PLGS were imported into DynamX 3.0 (Waters Corp.) to determine the deuterium uptake of samples. The following PLGS filtering parameters were applied: peptide quality thresholds of MS1 signal intensity greater than 5000, minimum products per amino acid of 0.3, maximum mass error of 1 ppm, retention time RSD of 0.5%, and PLGS score greater than 7. Following the automated peptide search and ion detection, each spectrum was manually inspected to ensure the corresponding m/z and isotopic distributions at various charge states were properly assigned to the appropriate peptic peptide. The uptake for all time points were averaged between three replicates. The percent relative deuterium uptake was calculated by dividing the relative deuterium uptake of each peptic peptide by its theoretical maximum uptake. Statistical significance for the differential HDX data is determined by t-test for each time point, and a 95% confidence limit for the uncertainty of the mean relative deuterium uptake was calculated to determine the threshold for significant uptake difference.

### Native mass spectrometry

Native MS analysis was performed after buffer exchanging proteins into the 150 mM ammonium acetate (AmAc) solution pH 6.8 using Amicon Ultra-0.5 Centrifugal Filter Units (Millipore Sigma). For the stoichiometry analysis of the UCH37•RPN13^DEUBAD^ and Nb-ncS1 complex, UCH37•RPN13^DEUBAD^ variants were premixed with Nb-ncS1 in the equimolar ratio and the solution was incubated on ice for 1 hour. For the analysis of Nb-ncS1 bound to UCH37•RPN13^DEUBAD^ -Ub-PA, the reaction between UCH37•RPN13^DEUBAD^ and Ub-PA was performed for 1 hour at 37°C prior mixing with Nb-ncS1. The samples were further diluted using 150 mM AmAc pH 6.8 to the final 10 uM concentration for the MS analysis. All native MS measurements were performed on a Synapt G2 HDMS (Waters Corp.) hybrid quadrupole/time-of-flight mass spectrometer equipped with a nanospray ion source. The following set of parameters was used: capillary voltage, 1.2 kV; sampling cone voltage, 40 V; extraction cone voltage 4 V. To enhance the transmission of the high-MW complexes, the backing pressure was increased up to 4 mBar. The MS data were analyzed with MassLynx 4.2 software (Waters Corp.); deconvolution of the mass spectra and calculation of the complex intact mass were performed using a UniDEC algorithm.

### AlphaFold 3 Modeling

Structural models of protein complexes were generated using AlphaFold 3 via the Google AlphaFold Server (https://deepmind.google/discover/alphafold), using the default multimer modeling settings. Amino acid sequences for UCH37, Ub, RPN13, and Nb-ncS1 were provided as input, reflecting the stoichiometry observed in the experimental system. No structural templates or experimental constraints were applied. Five models were generated per prediction, each revealing a similar, high-confidence inter-chain predicted TM (ipTM) score for the UCH37-Nb-ncS1 interface. Model confidence was assessed using ipTM and Predicted Aligned Error (PAE) plots. Figures were rendered in ChimeraX (v1.7) using the final AlphaFold 3 model.

### Protein crystallization and data processing

Individually purified UCH37 WT, RPN13 269-388, and cNb protein were incubated in a 1:2:2 ratio on ice for 4 hrs and subject to size exclusion chromatography using a HiLoad Superdex75 column (Cytiva) with 50 mM Tris pH 7.4, 100 mM NaCl, and 1 mM DTT as the mobile phase. Fractions of interest consisting of all three components of the complex were determined by SDS-PAGE Coomassie staining, pooled and concentrated to 25 mg/mL. The UCH37 WT, RPN13 269-388 and cNb complex was crystallized by sitting drop vapor diffusion at 21°C first identified in the commercial PEG Ion HT Screen (Hampton Research). Crystals grown in 0.1 M Sodium citrate tribasic dihydrate pH 5.6, 2% v/v Tacsimate pH 5.0, 16% w/v Polyethylene glycol 3,350 diffracted to 2.4 Å at the Advanced Photon Source (APS) at Argonne National Laboratory on the GM/CA 23-ID-B beamline. Data was processed using HKL3000 in the orthorhombic space group C222_1_.

### Structure determination and refinement

The structure of the UCH37 WT, RPN13 269-388, and Nb-cS1 complex was determined by molecular replacement using the program PHASER in the Phenix Suite. Input models used for molecular replacement were the heterodimer complex of UCH37 with RPN13 DEUBAD previously solved by Sixma and colleagues (PDB 4UEM) in addition to a homology model of the cNb generated by SWISS-MODEL with the primary sequence of the cNb and tertiary structure of the camelid Lag30 nanobody bound to eGFP (PDB 7SAI).

Model building and refinement was carried out using COOT and phenix refine where the current crystallographic *R* and *R_free_* are 0.235 and 0.271 respectively. Analysis of the Ramachandran plot indicated that 94.9% of residues fall within the most favored regions, 5.1% of residues fall within the allowed region, and none were observed in the disallowed region. Within the structure solution 65 of water molecules and 4 chloride ions are observed. While electron density is well resolved for structured regions, specific portions of the structure lacks depiction of a large number of residues. UCH37 WT is well resolved in regions 5-59, 74-149, 161-247, 253-312. All other regions, which includes about 55 residues, were not well resolved in the density. RPN13 269-388 was not well resolved in 31 residues which included residues 269-286, 357-359, and 385-388. The Nb-cS1 was not well resolved at positions 1-22, 48, 49, and 137. The final structure of the heterotrimer complex was validated using MolProbity and deposited in the Protein Data Bank under the accession code 9E7K.

### Ub-AMC assays

Ub-AMC assays were performed in black-well, 384-well plates (Corning). Mixtures containing UCH37•RPN13 (2 nM) with or without various concentrations of nanobodies in assay buffer (50 mM HEPES pH7.4, 50 mM NaCl, 0.5 mM TCEP) were incubated at 37°C for 15 min. 500 nM final concentration of Ub-AMC was then added to the appropriate wells, and hydrolysis was monitored continuously for 30 min at 37°C on a fluorescence plate reader (BioTek Synergy 2, λex = 360 nm, λem = 460 nm).

### Ub-Propargylamine (Ub-PA) labeling assay

2.5 μM of UCH37•RPN13 and indicated concentrations of Nb-cS1 or Nb-ncS1 in assay buffer (50 mM HEPES pH 7.4, 50 mM NaCl, 0.5 mM TCEP) were incubated at 37°C for 15 min. Ub-PA (2.5 μM) was then added and the reaction was incubated at 37°C for 30 min. Reactions were quenched by the addition of 6x Laemmli loading buffer. Samples were separated on a 15% SDS-PAGE gel and transferred on a PVDF membrane, stained with Streptavidin Alexa Fluor 647 (Thermo Fisher S32357) and imaged on the Odyssey CLx Imager (LI-COR Biosciences).

### Ub chain debranching assay

The debranching assays were performed as previously described (Deol et al., 2020), but with slight modifications. For the debranching assays, recombinant 1 μM UCH37•RPN13 with or without various concentrations of nanobodies were warmed to 37°C in DUB buffer for 15 minutes. Then 0.2 mg/ml HMW Ub chains or 10 μM branched Ub3 were added. Reactions were incubated for 1 hr at 37 °C and then quenched with 6x Laemmli loading buffer. Samples were separated on a 15% SDS-PAGE gel and either stained with SYPRO Ruby total protein stain or transferred on a PVDF membrane and blotted with mouse anti-ubiquitin antibody (Enzo Life Sciences, BML-PW0930-0100).

### Ub middle-down mass spectrometry (Ub MiD MS) analysis

K6/K48 HMW Ub chains treated with UCH37•RPN13 and nanobodies were quenched by freeze and thaw cycles. Lbpro* at final concentration of 15 μM was then added and incubated overnight to induce cleavage after R74 position of Ub^44^. The reaction was then quenched with 10% acetic acid (v/v) to precipitate other proteins except ubiquitin. The supernatant contains ubiquitin is dialyzed to water and reconstitute in 50% acetonitrile (v/v), and 0.1% formic acid (v/v). Samples were injected at a flow rate of 10 μl/min and ionized using a heated electrospray ionization (HESI-II) source before being analyzed using the Orbitrap Fusion Tribid mass spectrometer (Thermo Scientific). The resolving power of the mass analyzer on the spectrometer was set at 120000. All spectra were processed with in-house software (MASH Suite) using a signal-to-noise (S/N) threshold of 3 and a fit factor of 70% and then validated manually (Guner et al., 2014). Percentages correspond to the relative quantification values of the 11+ charge state for all three species: Ub1-74, 1xdiGly-Ub1-74, and 2xdiGly-Ub1-74.

### Kinetic analysis of Nb-ncS1 inhibition

Stock solutions of UCH37•RPN13, K6/K48Ub chains, and Nb-ncS1 were prepared in DUB buffer (50 mM HEPES pH 7.5, 50 mM NaCl, and 1 mM DTT).UCH37•RPN13 was kept constant at 0.5 μM with varied concentrations of K6/K48Ub chains and Nb-ncS1 as described in the figure. All reactions were performed at 37°C. Each sample along with a Ub and di-Ub standard was then separated on a 15% SDS-PAGE gel and stained with SYPRO Ruby. Gels were visualized on a Typhoon FLA 9500 (GE), and densitometry was performed using Image Studio. Initial velocities of Ub and di-Ub formation were converted to concentration per minute (µM/min). These values were then fit to the Michealis-Menten equation using nonlinear regression with competitive inhibition equation. Two biological replicates for each reaction were performed.

### Proximity ligation assay (PLA)

HEK293FT or HEK293FT UCH37KO cells were transfected with nanobodies as described above. 48 hr after transfection, cells were fixed with 4% paraformaldehyde and permeablized with 0.1% triton in PBS. The cells are then blocked with 3% BSA and incubated with rabbit anti-UCH37(Thermo Fisher, PA5-110544) or rabbit anti-UCHL3(Proteintech, 12384-1-AP) and mouse anti-HA (Cell Signaling Technology, 2367S) primary antibodies at 4 °C overnight. For PLA assay, the cells were incubated with Duolink In Situ PLA Probe Anti-Mouse MINUS and Anti-Rabbit PLUS, followed by ligation and amplification with Duolink In Situ Detection Reagents Red, and stained with DAPI. Cells were imaged with a Nikon TiE stand A1 Resonant Scanning Confocal microscope.

### Immunoprecipitation for western blotting

10 cm^2^ dishes of HEK293FT cells were grown and transfected with FLAG-tagged Nb in the 1436 pCDNA3 Flag HA vector (a gift from William Sellers, Addgene plasmids #10792). Transient transfections were performed using Xfect transfection reagent (Takara Bio) according to the manufacturer’s protocol. The harvested cells were lysed in 50 mM HEPES, pH 7.5, 50 mM NaCl, 5 mM MgCl_2_, 2 mM ATP, 1 mM DTT, and 10% glycerol before being incubated with anti-FLAG resin (Genscript) for 3 hours at 4 °C. The immunoprecipitated proteins were washed extensively with 50 mM HEPES, pH 7.5, 150 mM NaCl, 5 mM MgCl_2_, 2 mM ATP, 1 mM DTT, 0.05% IGEPAL prior to elution using 6x Laemmli loading buffer at 95 °C for 3 mins. Western blot analysis was performed using the following antibodies: rabbit anti-UCH37 (abcam, EPR4896), rabbit anti-ADRM1 (Cell Signaling Technology, D9Z1U), rabbit anti-PSMD14 (abcam, EPR4258), rabbit anti-PSMC1 (abcam, ab3317), mouse anti-PSMA2 (Cell Signaling Technology, 11864S) and rabbit anti-Flag (Cell Signaling Technology, 14793S) at 1:1000 dilution. IRDye® 680RD goat anti-rabbit and IRDye® 800CW goat anti-mouse (LI-COR Biosciences) were used as the secondary antibodies at 1:15000 dilution.

### Immunoprecipitation for mass-spectrometry

15 cm^2^ dishes of HEK293FT cells were grown and transfected with HaloTag nanobodies as described above. The harvested cells were lysed and immunoprecipitated similarly as described above. The immunoprecipitated proteins were washed extensively with 50 mM HEPES, pH 7.5, 150 mM NaCl, 5 mM MgCl_2_, 2 mM ATP, and 1 mM DTT. The bound proteins were reduced with 10 mM DTT at 37 °C for 1 hr and alkylated using 25 mM iodoacetamide (Sigma-Aldrich) in 25 mM NH_4_CO_3_ at room temperature for 1 hr in the dark before being subjected to on-bead trypsin digestion using sequencing grade modified trypsin (Promega) for overnight at 37 °C. Tryptic digests were desalted with Sep-Pak C18 cartridge (Waters), quantified by Pierce Quantitative Colorimetric Peptide Assay, and then subjected to TMT labeling. 10 μL of each sample (20-40 μg) peptides in 100 mM TEAB were added with 5 μL of 20 ng/μL 10-plex TMT reagents (Thermo Scientific) dissolved in anhydrous acetonitrile. Following incubation at RT for 1 hr, the labeling reaction was quenched with hydroxylamine to a final concentration of 0.4 % (v/v) for 15 min, and the equal volume of TMT labeled samples were pooled together. Peptides were desalted with Sep-Pak C18 cartridge (Waters) and separated by a homemade fused silica capillary column (75 μm x 150 mm) packed with C-18 resin (120A, 1.9 μm, Dr. Maisch HPLC GmbH) with the EASY-nLC 1000 nano-HPLC system (ThermoFisher Scientific), which was coupled with the Orbitrap Fusion Tribrid mass spectrometer.

The scan sequence began with an MS1 spectrum (Orbitrap analysis; resolution 120,000; mass range 375–1500 m/z; automatic gain control [AGC] target 5 × 10^5^; maximum injection time 50ms). MS2 analysis consisted of CID with 35% energy (AGC 5 × 10^3^; isolation window 1.2 Th; maximum injection time 50 ms; activation time 10 ms). Monoisotopic peak assignment was used, and previously interrogated precursors were excluded using a dynamic window (15 s ± 10 ppm), and dependent scans were performed on a single charge state per precursor. Following the acquisition of each MS2 spectrum, a synchronous-precursor-selection (SPS) MS3 scan was collected on the top 10 most intense ions in the MS2 spectrum. MS3 precursors were fragmented by high energy CID with 65% energy and analyzed using the Orbitrap (AGC 5 × 10^4^ maximum injection time 86 ms, resolution was 50,000 at 2 Th isolation window).

Raw files were processed using Proteome Discoverer 2.5 (Thermo Fisher Scientific) with the SEQUEST HT search engine. The workflow was configured for SPS MS3–based TMT reporter ion quantification, using MS2 (CID) for peptide identification and MS3 (HCD) for quantification. Spectra were searched against the UniProt Homo sapiens database (release: December 14, 2022). Search parameters included trypsin digestion with up to two missed cleavages, a precursor mass tolerance of 10 ppm, and a fragment mass tolerance of 0.6 Da. Carbamidomethylation of cysteine (+57.021 Da) and TMT 10-plex labeling of peptide N-termini and lysine residues (+229.163 Da) were specified as static modifications. Methionine oxidation (+15.995 Da) and N-terminal acetylation (+42.011 Da) were included as dynamic modifications. Peptide-spectrum matches were filtered to a 1% false discovery rate (FDR) using the Percolator algorithm. For statistical analysis of IP-MS enrichment, a two-sample t-test was performed and proteins were considered significantly enriched at p < 0.05 and log2 fold change > 1.

## DATA AVAILABILITY

The structural coordinates for Nb-cS1 in complex with UCH37•RPN13^DEUBAD^ have been deposited in RCSB Protein Data Bank (PDB) under accession code 9E7K. Source data are provided within this paper (Source data.zip). The HDX-MS data have been deposited in MassIVE under the accession codes MSV000098855, MSV000098853, and MSV000098852. Proteomics data corresponding to the IP-TMT-MS of Nb-ncS1 and Nb-cS1 transfected cells have been deposited under the accession code MSV000096425. Additional data can be obtained from the corresponding author upon request.

## ACKNOWLEDGEMENTS

We thank Dr. Steve Eyles (UMass Amherst) for assistance with high-resolution MS at UMass Amherst Mass Spectrometry Core Facility (RRID:SCR_019063), Dr. Amy S. Burnside (UMass Amherst) for assistance with flow cytometry, and Dr. Ravi Ranjan for the NGS experiment at the UMass Amherst Genomics Resource Laboratory.

MST, SPR and SEC-MALS data were acquired in the UMass Amherst Biophysical Characterization Core Facility (RRID:SCR_022357). The PLA data was gathered at the UMass Amherst Light Microscopy Facility. This work was funded by the National Institutes of Health (NIH) (R35GM149532 to E.R.S. and R01GM126296 to C.D.), an NIH Chemistry and Biology Training Grant (T32GM139789 to J.D.), and an NIH Ruth L. Kirschstein NRSA (F31CA275390 to R.P.). The data described herein were acquired on an Orbitrap Fusion mass spectrometer funded by NIH grant 1S10OD010645-01A1.

## AUTHOR CONTRIBUTIONS

ERS: Conceived the project, conceptualization, funding acquisition, supervision, and data analysis. YL, LHC, ES, JD, MS, and ANRD: Generated and characterized the nanobodies. RP and CD: Solved and analyzed the crystal structure. YL, ES and ERS wrote the manuscript.

