## Supplemental Data for "Site-Specific Nanobody Inhibitors of the Proteasomal Deubiquitinase UCH37"

### SUPPLEMENTARY DATA

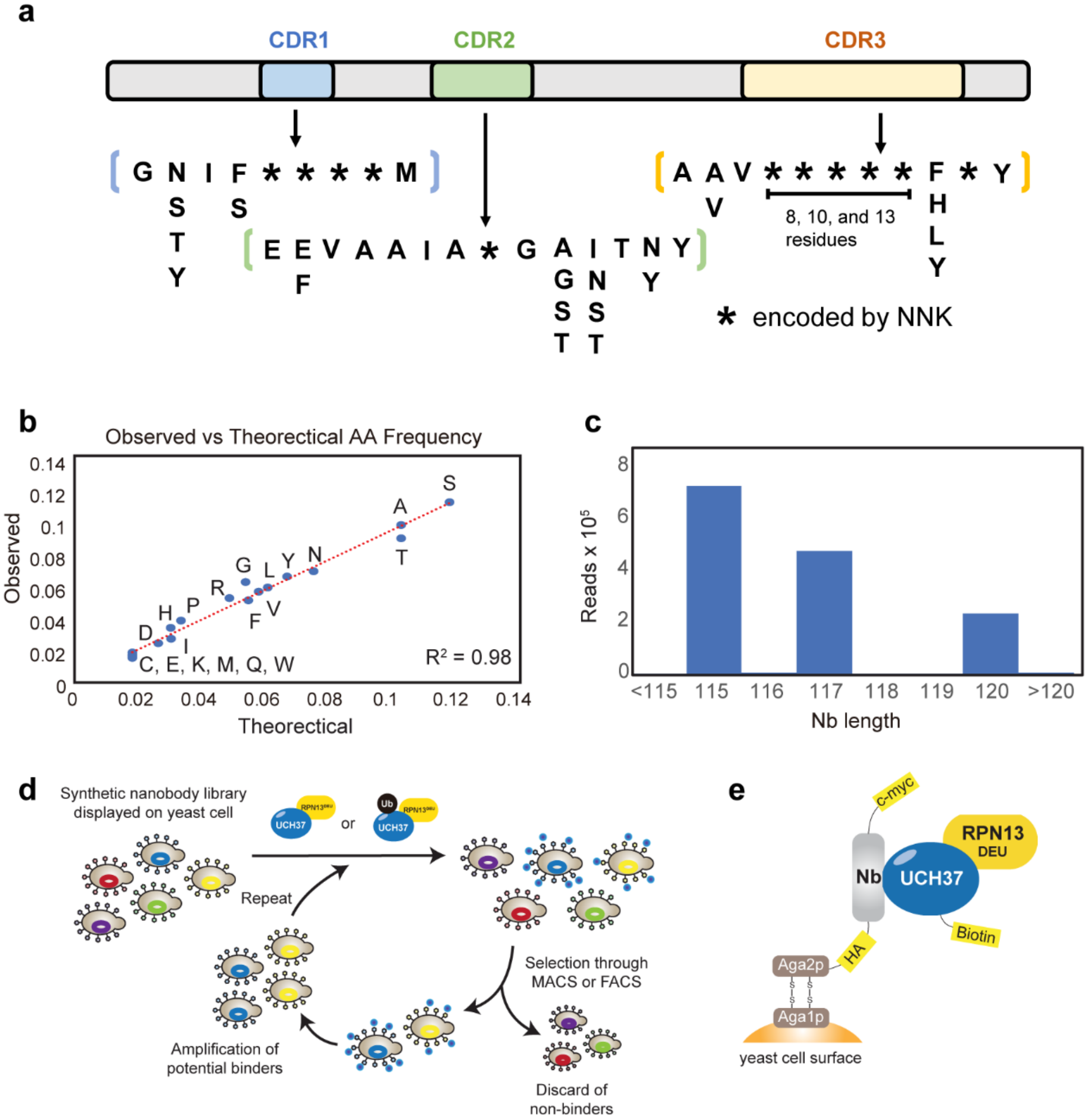


**Supplementary Figure 1: Construction and NGS Analysis of the Synthetic Nanobody Library**

(a) General scheme of the synthetic nanobody highlights fixed framework regions (shown in gray), while the CDR loops: CDR1 (blue), CDR2 (green), and CDR3 (orange) were designed with variations. Partial randomization in these loops was introduced by PCR using primers with different degenerate codons. Highly variable regions were introduced by PCR using primers with NNK codons, marked by asterisks (*).

(b) The amino acid frequencies at the diversified positions within the CDRs. The results demonstrated that the distribution of amino acids in the library closely aligned with the target values.

(c) Bar plot showing the distribution of the ensemble of nanobodies based on their length.

(d) General scheme of the nanobody selection process. Yeast displaying nanobodies with affinity to antigen are shown being isolated via MACS or FACS, amplified, and undergoing iterative rounds of selection

(e) General scheme of the yeast display platform. The nanobody is shown with an N-terminal HA tag and a C-terminal c-myc tag, and fused to the C-terminal of Aga2p which tethers the nanobody to the yeast cell wall.

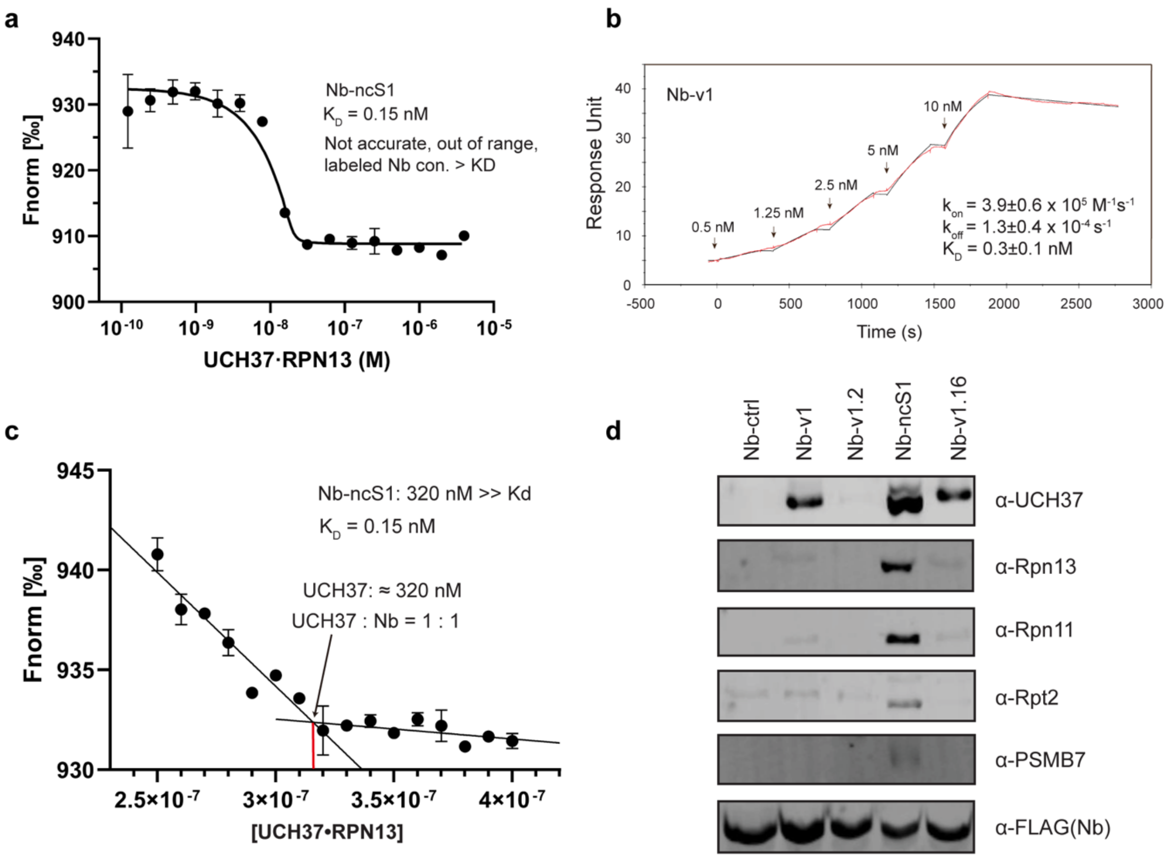


**Supplementary Figure 2: Biophysical Characterization of Nb-v1 and Nb-ncS1**

(a) Microscale thermophoresis (MST) experiments showing affinity between Nb-ncS1 and UCH37•RPN13.The measured $K_D$ value (0.15 nM) is far below the concentration of the fluorescently labeled Nb-ncS1 (20 nM) and the $K_D$ is outside the optimal range for the instrument.

(b) Single-cycle surface plasmon resonance (SPR) experiment showing the interaction between Nb-v1 and UCH37. Experimental data is in red, and the curve fit is in black.

(c) Microscale thermophoresis (MST) experiments showing stoichiometry between Nb-ncS1 and UCH37•RPN13. Fluorescently labeled Nb-ncS1 is used at a concentration much higher than Kd (320 nM vs 0.13 nM) to ensure the UCH37•RPN13 titrated in is fully occupied and produces a signal. The kink point is where all Nb-ncS1 is bound by UCH37•RPN13

(d) Western blot analysis of anti-FLAG pull-downs with HEK293FT cell lysates transfected with FLAG-Halo-tagged nanobodies. Samples were separated by SDS–PAGE and immunoblotted against UCH37 along with other proteasome subunits. Nb-v1.2 and Nb-v1.16 are two other mutants of Nb-v1.

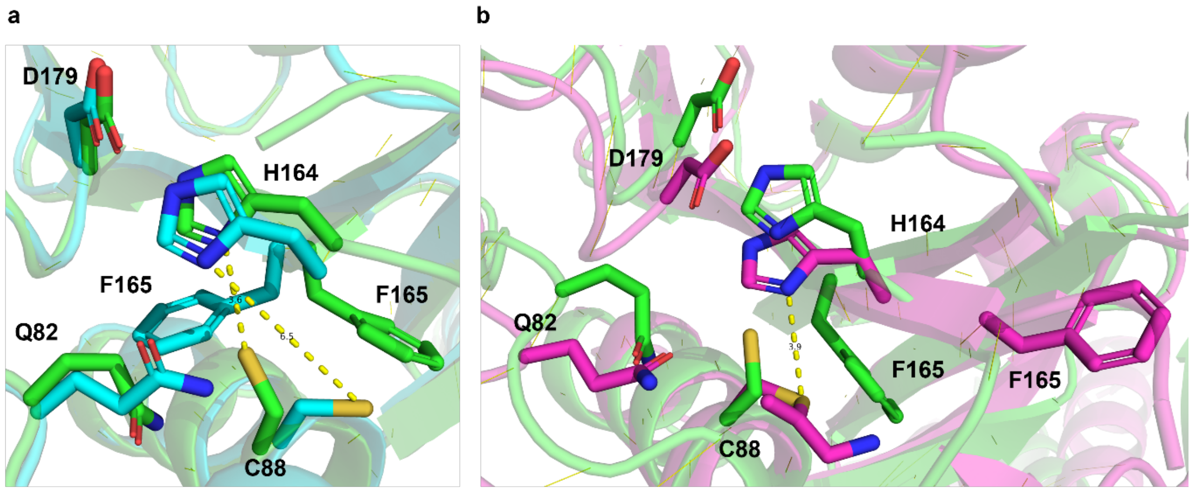


**Supplementary Figure 3: Nb-cS1-Induced Conformational Change of UCH37 Active Site**
Structural comparison of the Nb-cS1 with UCH37•RPN13$^{DEUBAD}$ complex (green) with the UCH37 structures from the apo state (PDB 4UEM, cyan) and Ub-bound state (PDB 4UEL, purple), focused on the active site of UCH37. Key residues involved in the active site—D179, H164, F165, Q82, and C88—are highlighted. In the apo state (4UEM and PDB 3RII), the catalytic triad (C88, H164, D179) is misaligned. Monoubiquitin binding, as shown in the Ub-bound structure (4UEL), induces a conformational change that aligns C88 with H164, promoting a productive conformation. Our Nb-cS1-bound UCH37 structure also promotes alignment of the catalytic triad into a productive conformation while occupying the cS1 site, suggesting the mechanism of inhibition might not be dominated through active site rearrangement.

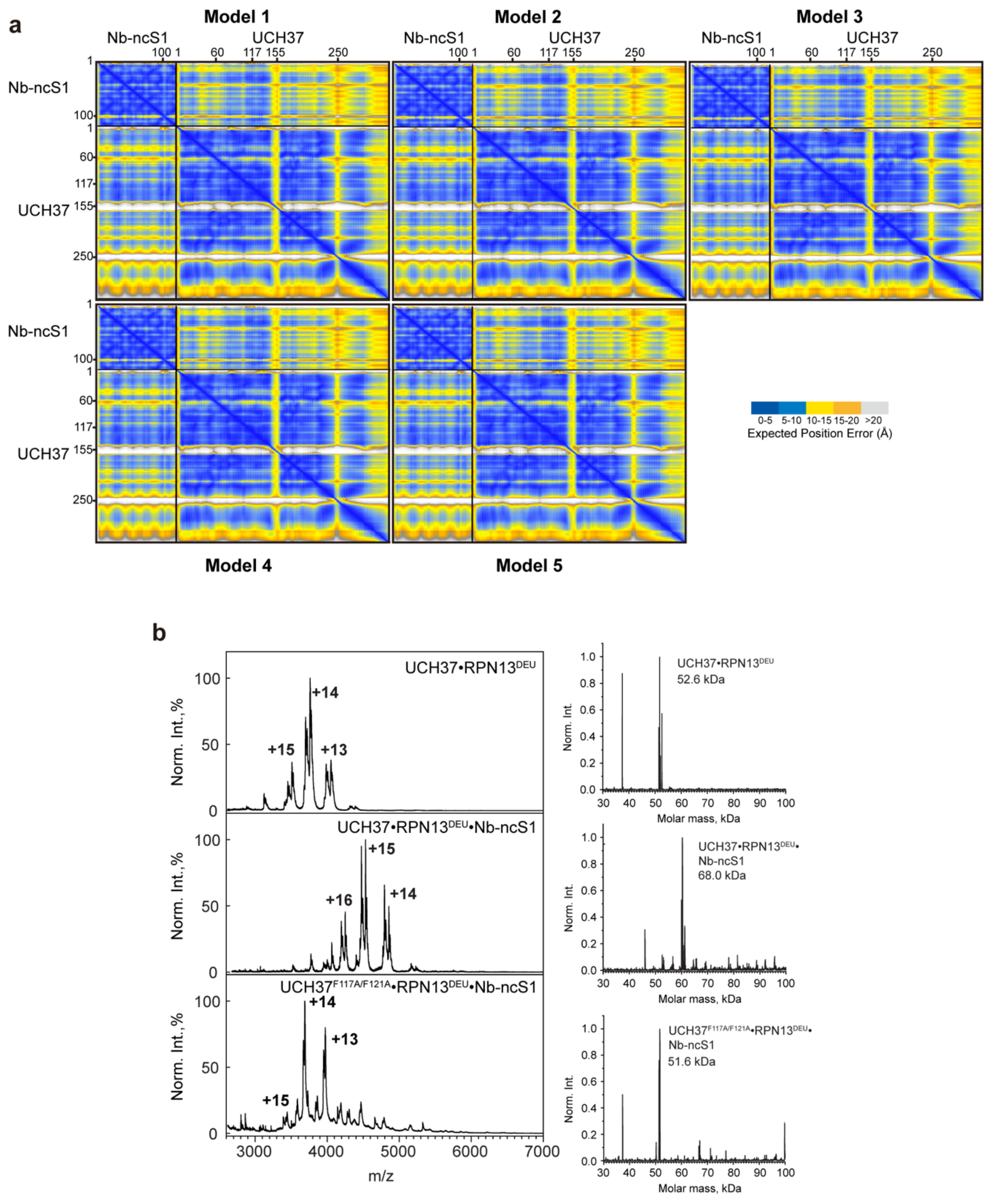


**Supplementary Figure 4: Interaction between Nb-ncS1 and UCH37 is Predicted to Occur on the Backside of UCH37.**

(a) Predicted Alignment Error (PAE) plots for Nb-ncS1 bound to UCH37. The x- and y- axes correspond to residue indices for Nb-ncS1 and UCH37. Colors indicate the expected positional error (in Å) at residue *x* when the predicted and actual structures are aligned at residue *y*, with blue representing high-confidence alignment (low error) and lighter colors to orange/gray indicating lower confidence (higher error).

(b) Native ESI-MS spectra of free UCH37•RPN13$^{DEUBAD}$ (top panel), Nb-ncS1-bound UCH37•RPN13$^{DEUBAD}$ (middle panel), and Nb-ncS1 bound to Ub-UCH37$^{F117A/F121A}$•RPN13$^{DEUBAD}$ (bottom panel). Additional peaks observed in the +13 to +15 charge states for free UCH37·RPN13$^{DEUBAD}$ and in the +14 to +16 charge states for the complex correspond to N-/C-terminal truncations in the DEUBAD domain of RPN13. Deconvoluted spectra corresponding to each raw trace are shown to the right.

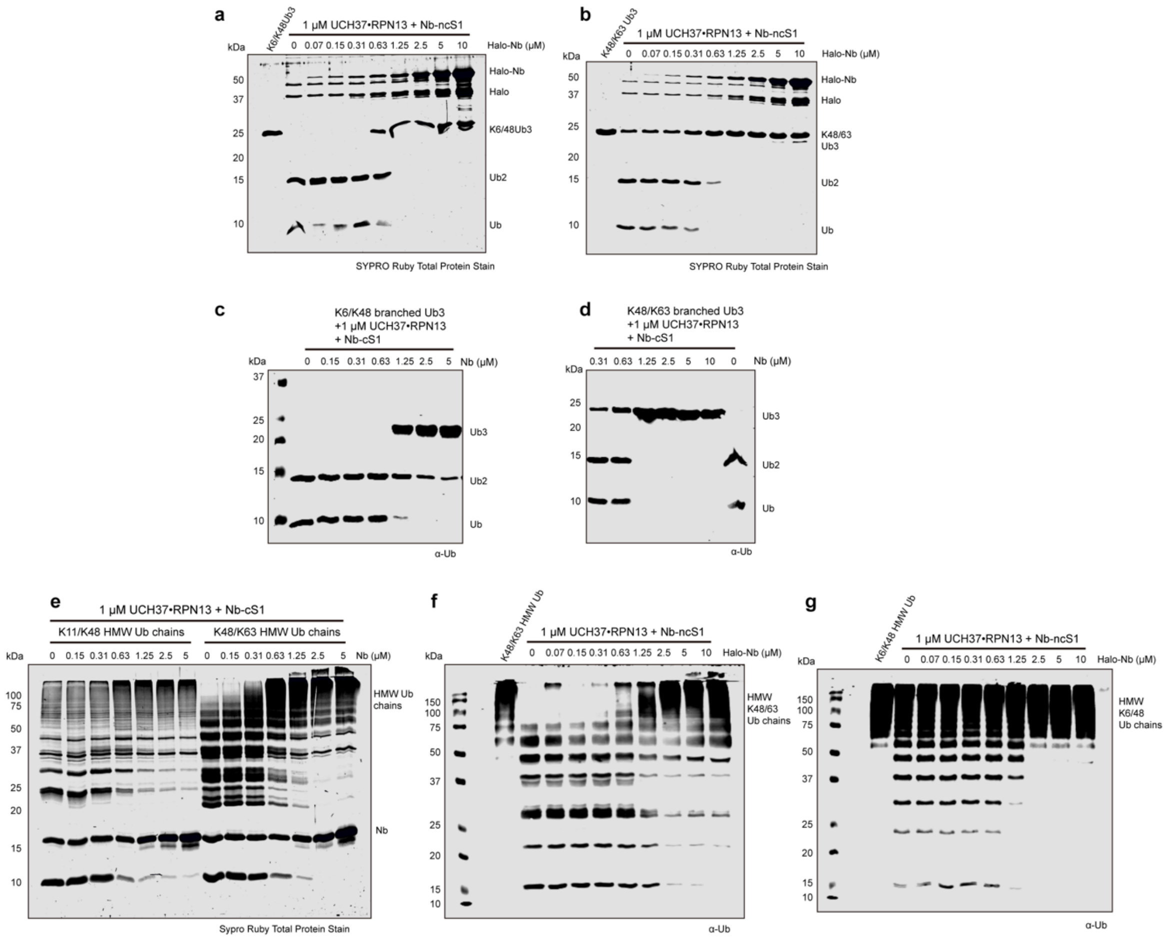


**Supplementary Figure 5: Nb-ncS1 and Nb-cS1 Both Inhibit the K48 Debranching Activity of UCH37**

(a-b) SDS-PAGE analysis of the cleavage of K6/K48 (a) or K48/K63 (b) branched trimer with 1 µM of UCH37•RPN13 co-incubated with different concentrations of Halo-tagged Nb-ncS1.

(c-d) Western blot analysis of the cleavage of K6/K48 (c) or K48/K63 (d) branched trimer with 1 µM of UCH37•RPN13 co-incubated with different concentrations of Nb-cS1.

(e) SDS-PAGE analysis of the cleavage of K11/K48 or K48/K63 high-molecular weight (HMW) Ub chains with 1 µM of UCH37•RPN13 co-incubated with different concentrations of Nb-cS1.

(f) Western blot analysis of the cleavage of K48/K63 high-molecular weight (HMW) Ub chains with 1 µM of UCH37•RPN13 co-incubated with different concentrations of Halo tagged Nb-ncS1.

(g) Western blot analysis of the cleavage of K6/K48 high-molecular weight (HMW) Ub chains with 1 µM of UCH37•RPN13 co-incubated with different concentrations of Halo tagged Nb-ncS1.

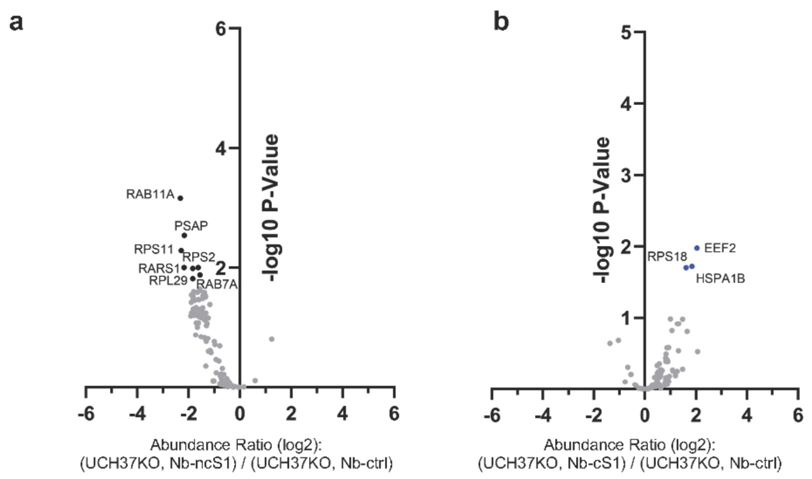


**Supplementary Figure 6: Nb-ncS1 and Nb-cS1 Did Not Pull-Down Proteasome in UCH37KO Cells**

(a-b) Volcano plot from IP-TMT-MS data comparing proteins enriched by Halo-tagged Nb-ncS1 (a) or Halo-tagged Nb-cS1 (b) versus the Halo-tagged control nanobody (Nb-ctrl).

| Primers | Sequence (5'-3') |
|---|---|
| a1_f | GCTAGCCAGGTGCAGCTGCAGGAAAGCGGCGGCCTGG TGCAGGCGGGC |
| a2_r | GCCGCTCGCCGCGCAGCTCAGGCGCAGGCTGCCGCCCGCC TGCAC |
| b3_f | CGCGGCGAGCGGCWMTATTTYTNNKNNKNNKNNKATGGGCT GGTAT |
| b4_r | AATAGYAGCAACAAGTTCGCGTTCTTTGCCCGGCGCCTGGCG ATACCAGCCCAT |
| c5_f | GTTGCTRCTATTRVTNNKGGTRSTANTACCWATTATGCGGATA GCGTGAAAGGCCGCTTT |
| c6_r | TTCATCTGCAGATACACGGTGTTTTTCGCGTTATCGCGGCTAA TGGTAAAGCGGCCTTT |
| d7_f | TCTGCAGATGAACAGCCTGAAACCGGAAGATACCGCGGTGTA TTATTGCGCGG |
| d8_short_r | CCCCAATATGCAWRMNNMNNMNNMNNMNNARCCGCGCAATA AT |
| d8_medium_r | CCCCAATATGCAWRMNNMNNMNNMNNMNNMNNMNNARCCG CGCAATAAT |
| d8_long_r | CCCCAATATGCAWRMNNMNNMNNMNNMNNMNNMNNMNNMN NMNNARCCGCGCAATAAT |
| e9_f | GCATATTGGGGCCAGGGCACCCAGGTGACCGTGAGCAGCGG TGGTGGTGGTAGC |
| e10_r | GTCCCCGAAGTTCAGACCGGTGCTACCACCACCACCGCTGCT CACGGT |
| pCT_nb_PstI_f | GGTGGTGGTTCTGCTAGCCAGGTGCAGCTGCAGGAA |
| pCT_nb_Bam HI_r | TTACAAGTCCTCTTCAGAAATAAGCTTTTGTTCGGATCCGCTAC CACCACCGCT |

**Supplementary Table 1: Primers Used in the Generation of the Nanobody Library**

| Nb variants | Sequence |
|---|---|
| Nb-v1 | QVQLQESGGGLVQAGGSLRLSCAASGYIFRSYRMGWYRQAPGKERELVAAIGLGGNTYYADSVKGRFTISRDNAKNTVYLQMNSLKPEDTAVYYCAALHGKPPAFSPFAYWGQGTQVTVSS |
| Nb-ncS1 | RVQLQESGGGLVQAGGSLRLSCAASGYIFRSYRMGWYRQAPGKERELVAAIGLGGNTYYADSVKGRFTISRDNAKNTVYQQMNSLKPEDTAVYYCAALYGKPPAFSPFPYWGQGTQVTVSS |
| Nb-v2 | QVQLQESGGGLVQAGGSLRLSCAASGYISRPRTMGWYRQAPGKERELVATISNGAITNYADSVKGRFTISRDNAKNTVYLQMNSLKPEDTAVYYCAAGKNIWHAYWGQGTQVTVSS |
| Nb-cS1 | QVQLQESGGGLVQAGGSLRLSCAASGYISRPRTMGWYRQAPGKEREFVAIISNGAITNYADSVKGRFTISRDNAKNTVYLQMNSLKPEDTAVYYCAAGRNIWHAYWGQGTQVTVSS |
| Nb-ctrl | QVQLQESGGGLVQAGGSLRLSCAASGSISKWGLMGWYRQAPGKERELVAAIDSGANTYYADRVKGRFTISRDNAKNTVYLQMNSLKPEDTAVYYCAVVAIRRSDHADWGQGTQVTVSS |

**Supplementary Table 2: Sequences of the Nanobodies**

| UCH37 – RPN13 DEU Nb-cS1 | |
|---|---|
| **Data Collection and Processing** | |
| Beamline | APS GM/CA 23-ID-B |
| Resolution range (Å) | 37.3 – 2.4 (2.49-2.40) |
| Space group | $C222_1$ |
| Cell dimensions | |
| a, b, c (Å) | 45.56, 178.63, 190.73 |
| α, β, γ (˚) | 90, 90, 90 |
| Total reflections | 99575 (10358) |
| Unique reflections | 29105 (2823) |
| Redundancy (%) | 3.4 (3.7) |
| Completeness (%) | 94.58 (94.16) |
| $R_{merge}$ | 0.066 (0.831) |
| $R_{meas}$ | 0.079 (0.978) |
| $R_{pim}$ | 0.043 (0.508) |
| I/σ(I) | 14.72 (1.62) |
| CC1/2 | 0.998 (0.532) |
| Wilson B (Å$^2$) | 68.85 |
| **Refinement** | |
| Complex/A.S.U., Complex Ratio | 1, 1:1:1 |
| Resolution (Å) | 2.40 |
| Rwork / Rfree | 0.235 / 0.271 |
| No. nonhydrogen atoms | 3770 |
| Protein | 3701 |
| Water | 65 |
| Chloride ions | 4 |
| B factors (Å$^2$) | 87.58 |
| Protein | 87.74 |
| Water | 77.99 |
| Chloride ions | 95.50 |
| R.m.s.d | |
| Bond lengths (Å) | 0.003 |
| Bond angles (˚) | 0.59 |
| Ramachandran (favored/allowed/outliers) | 94.9 %, 5.1 %, 0 % |
| Clash score | 8.01 |

**Supplementary Table 3:** Data collection, processing, and refinement statistics for the X-ray crystal structure of UCH37 RPN13 DEUBAD complex bound to Nb-cS1. Data is deposited in the Protein Data Bank under the accession code 9E7K.
